# Metabolite discovery through global annotation of untargeted metabolomics data

**DOI:** 10.1101/2021.01.06.425569

**Authors:** Li Chen, Wenyun Lu, Lin Wang, Xi Xing, Ziyang Chen, Xin Teng, Xianfeng Zeng, Antonio D. Muscarella, Yihui Shen, Alexis Cowan, Melanie R. McReynolds, Brandon Kennedy, Ashley M. Lato, Shawn R. Campagna, Mona Singh, Joshua Rabinowitz

## Abstract

Liquid chromatography-high resolution mass spectrometry (LC-MS)-based metabolomics aims to identify and quantitate all metabolites, but most LC-MS peaks remain unidentified. Here, we present a global network optimization approach, NetID, to annotate untargeted LC-MS metabolomics data. The approach aims to generate, for all experimentally observed ion peaks, annotations that match the measured masses, retention times, and (when available) MS/MS fragmentation patterns. Peaks are connected based on mass differences reflecting adducting, fragmentation, isotopes, or feasible biochemical transformations. Global optimization generates a single network linking most observed ion peaks, enhances peak assignment accuracy, and produces chemically-informative peak-peak relationships, including for peaks lacking MS/MS spectra. Applying this approach to yeast and mouse data, we identified five novel metabolites (thiamine derivatives and N-glucosyl-taurine). Isotope tracer studies indicate active flux through these metabolites. Thus, NetID applies existing metabolomic knowledge and global optimization to annotate untargeted metabolomics data, revealing novel metabolites.

## Introduction

Metabolomics provides a snapshot of the concentrations of detectable small-molecule in a biological system. In so doing, it reflects the integrated impact of genetics and environment on metabolism. One important role of metabolomics is annotating previously unknown or underappreciated metabolites. For example, metabolomics facilitated identification of 2-hydroxyglutarate as an oncometabolite, eventually leading to the development of inhibitors of 2-hydroxyglutarate synthesis as anticancer agents^1,2^. Metabolomics also contributed to the identification of a diversity of natural products^3,4^ and disease biomarkers^5^.

A common experimental strategy in metabolomics is liquid chromatography-high resolution mass spectrometry (LC-MS). LC-MS metabolomics measures thousands of ion peaks, of which hundreds are associated with known metabolites. A much greater number of peaks, however, still remain unannotated. A common approach to peak annotation is to compare exact mass and either retention time or MS/MS (MS2) fragmentation pattern to authenticated standards. To facilitate such comparisons, extensive molecular structural databases (Pubchem^6^, HMDB^7^, KEGG^8^, ChemSpider^9^), MS2 spectral databases (METLIN^10^, GNPS^11^, MassBank^12–14^ and NIST^15^) and software (e.g. XCMS^16,17^, GNPS^11^, SIRIUS^18^ and MS-DIAL^19^) have been developed. Peaks can also arise from mass spectrometry phenomena, such as adducts, fragments or isotopes of metabolites^20–24^. Such peaks seem to account for at least half of non-background LC-MS features^25–27^. Despite this progress, a great number of unknown peaks remain, and figuring out their identities is a primary challenge in the field.

Network analysis, capitalizing on peak-peak relationships to increase annotation scope and accuracy, has been broadly used in metabolomics data annotation. Workflows employing the concept of molecular connectivity have been used to build networks (e.g., GNPS^28–30^, MetDNA^31^, CliqueMS^32^ and others^33–35^). Ions connected by either biochemistry or mass spectrometry phenomena often share MS2 fragmentation pattern similarity. While distinct metabolites typically separate chromatographically, ions connected through mass spectrometry phenomena co-elute.

Discovery of novel metabolites involves generation of candidate molecular formulae beyond those in current databases. This can be achieved by modifying formulae of known metabolites using characteristic biochemically feasible atom transformations that match observed MS1 mass differences^35,36^ (e.g. 2.016 for 2H). Such approaches can be combined with clustering metabolites based on similar MS2 fragmentation patterns in a molecular network, as demonstrated in GNPS and other works^28–31,37^. In a cluster of connected peaks, one known metabolite peak can help to annotate its neighbors, facilitating unknown discovery. One state-of-the-art database-independent method to generate novel candidate molecular formulae (SIRIUS 4.0) combines high-resolution m/z, natural isotope abundances, and MS2 spectral analyses^18^.

Existing methods generally focus on annotation of either individual peaks or a subnetwork of peaks. All peaks can be sequentially assessed, but individual annotations do not make full use of all available information regarding other peaks in the network. Global network optimization methods, not dealing with peak annotation one-by-one, but instead all at once to take full advantage of the entire available information, hold potential. For example, a Gibbs-sampling statistics approach has been used to incorporate biochemical connections, isotope patterns, and adduct relationships for probabilistic annotations of all peaks in a metabolomics experiment^38–40^. ZODIAC combines a similar approach and the SIRIUS algorithm^41^. Such Gibbs-sampling approaches allow network connections to shift the probabilities of candidate peak annotations, leading to substantially better annotations results^41^. While capable of assessing the probability of all individual candidate annotations in light of their neighbors, broadly defined, such approaches do not guarantee selection of the optimal overall network as a whole.

An alternative computational strategy focusing on optimizing the entire network, as opposed to statistical annotation of individual peaks, is integer linear programming. Given a scoring function, this computational optimization strategy determines the single best annotation for all peaks collectively. Global optimization of large networks of many thousands of peaks is feasible due to linear formulation of the problem, which renders the approach computationally efficient.

Integer linear programming optimization has, to our knowledge, not been previously applied in the context of molecular network analysis. To this end, we present an efficient stand-alone algorithm “NetID”. The algorithm optimizes a network of mass spectrometry peak connections based on MS1 mass differences corresponding to gain or loss of relevant chemical moieties and MS2 spectral similarity, in a manner that differentiates biochemical connections from those based on mass spectrometry phenomena, and that incorporates literature data on known metabolites and their retention times. We applied this integer linear programming optimization approach to untargeted metabolomics data from both Baker’s yeast and mouse liver. The global optimization step enforces a single formula assignment for each experimentally observed ion peak, increasing annotation accuracy as estimated by a target–decoy strategy^42^. Through these efforts, we provide likely formulae for several hundred putative novel metabolites and confirm the identities of five new metabolites.

## Results

### NetID algorithm

NetID involves three computational steps: candidate annotation, scoring, and network optimization (Figure 1). The workflow starts with a peak table that contains a list of peak m/z, RT, intensity, and (when available) associated MS2 spectra, with background peaks removed by comparing to a process blank sample. Each peak defines a node in the network. In the candidate annotation phase, we match every experimentally measured node m/z to formulae in the selected metabolomics database (e.g. HMDB). Peaks matching to database formula within 10 ppm are assigned as seed nodes with candidate seed formulae, from which we extend edges to build the network.

**Figure 1.**
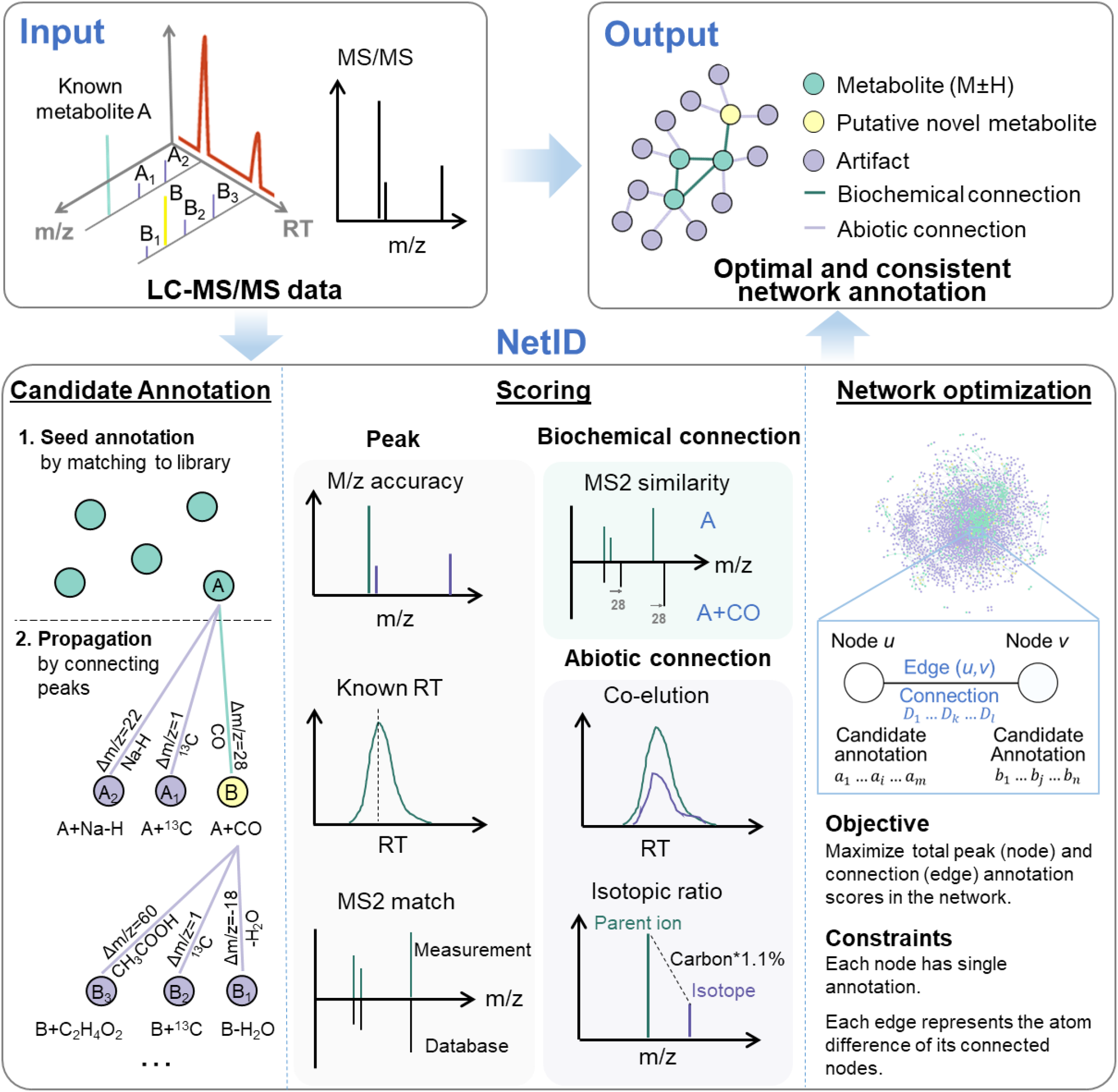
A global network optimization approach for untargeted metabolomics data annotation (NetID). The input data are LC-MS peaks with m/z, retention times, intensities and optional MS2 spectra. The output is a molecular network with peaks (nodes) assigned with unique formulae and connected by edges reflecting atom differences arising either through metabolism (biochemical connection) or mass spectrometry phenomenon (abiotic connection). Peaks are classified as “metabolite” (M+H or M-H peak of formula found in selected metabolomics database, e.g. HMDB), “putative novel metabolite” (formula not found in database but with biochemical connection to a metabolite), or “artifact” (only abiotic connection to a metabolite). NetID algorithm involves three steps. Candidate annotation first matches peaks to database formulae. These seed annotations are then extended through edges to cover most nodes, with the majority of nodes receiving multiple formula annotations. Each node and edge annotation are then scored based on match to known masses, retention times, and MS/MS fragmentation patterns. Global network optimization maximizes sum of node scores and edge scores, while enforcing a unique formula for each node and unique transformation relationship for each edge.

Edges connect two nodes via gain or loss of specific chemical moieties (atoms). The atom differences can occur either due to metabolism (biochemical connection) or due to mass spectrometry phenomena (abiotic connections). For example, a difference of H_2_ suggests an oxidation/reduction relationship and defines a biochemical edge. A difference of Na-H suggests sodium adducting and is a type of abiotic edge (adduct edge). Other atom differences define other types of abiotic connections (isotope or fragment edges). Most atom differences are specific to biochemical, adduct, isotope, or fragment edges, but a few occur in multiple categories. For example, H_2_O loss can be either biochemical (enzymatic dehydration) or abiotic (in-source water loss). By integrating literature and in-house data, we assembled a list of 25 biochemical atom differences and 59 abiotic atom differences which together define all connections in the network (Supplementary Table 1, 2). Using these lists, we make candidate edge annotations such that (i) ∆m/z between the connected nodes matches the atom mass difference and (ii) only co-eluting peaks are connected by abiotic edges. Through the edge extension process starting from the seed nodes, candidate formulae are assigned to nodes outside the initial seeds. A few rounds of edge extension suffice to give thorough coverage (see Methods). Due to finite mass measurement precision, a single node (including a seed node) may be assigned multiple contradictory candidate formulae, which are resolved at the following scoring and optimization step.

NetID then scores every candidate node and edge annotation. Candidate node annotations are scored based on precision of m/z match to the molecular formula and (when the relevant information is available) precision of retention time match to known metabolite retention time and quality of MS2 spectra match to database structure. In addition, there is a bonus for matching to formula in HMDB and a penalty for unlikely formulae (e.g. containing an uncommon elemental ratio or extreme number of ring and double bond equivalents)^43^. Biochemical edges receive a positive score for MS2 spectra similarity between the connected nodes. Abiotic edges are scored based on precision of co-elution with the parent metabolite, connection type (adduct, isotope, etc.), and features specific to the connection type, such as expected natural abundance for isotope peaks (see Methods and Supplementary Note 2). The overall impact is to assign high scores to those candidate annotations that effectively align the experimentally observed ion peaks with prior metabolomics knowledge.

With a score assigned for each candidate node and edge annotation, we formulate the global network optimization problem as that of maximizing the network score with linear constraints that each node and edge has a single annotation and that they are consistent (e.g. peaks connected by H2 edge must have formula differing by 2H). Such optimization is readily performed by linear programing with a typical runtime of minutes to hours on a personal computer, and results in an optimal and consistent network annotation.

### Global network optimization

As an example of the utility of global network optimization, where all peaks and connections are simultaneously considered to enhance annotation accuracy, we present an example network containing five peaks (Figure 2A). We first match experimental measurements to the database, assigning node *a* and node *b* as seed nodes adenosine monophosphate (AMP, C_10_H_14_N_5_O_7_P) and adenosine (C_10_H_13_N_5_O_4_), respectively. We also identify five possible connections between the five nodes. Two alternative networks are generated by extending from seed assignments. In the left network, node *c* is annotated as adenosine HCl adduct (C_10_ClH_14_N_5_O_4_), whereas in the right network, node *c* is (mis)annotated as a putative novel metabolite (C_9_H_14_N_5_O_5_P) resulting from CO_2_ loss from AMP. Node *d* is ^13^C isotope of node *c* in both networks. Node *e* is annotated as ^37^Cl isotope of node *c* in the left network, and is unannotated in the right network because there is no Cl atom in the parent molecule.

**Figure 2.**
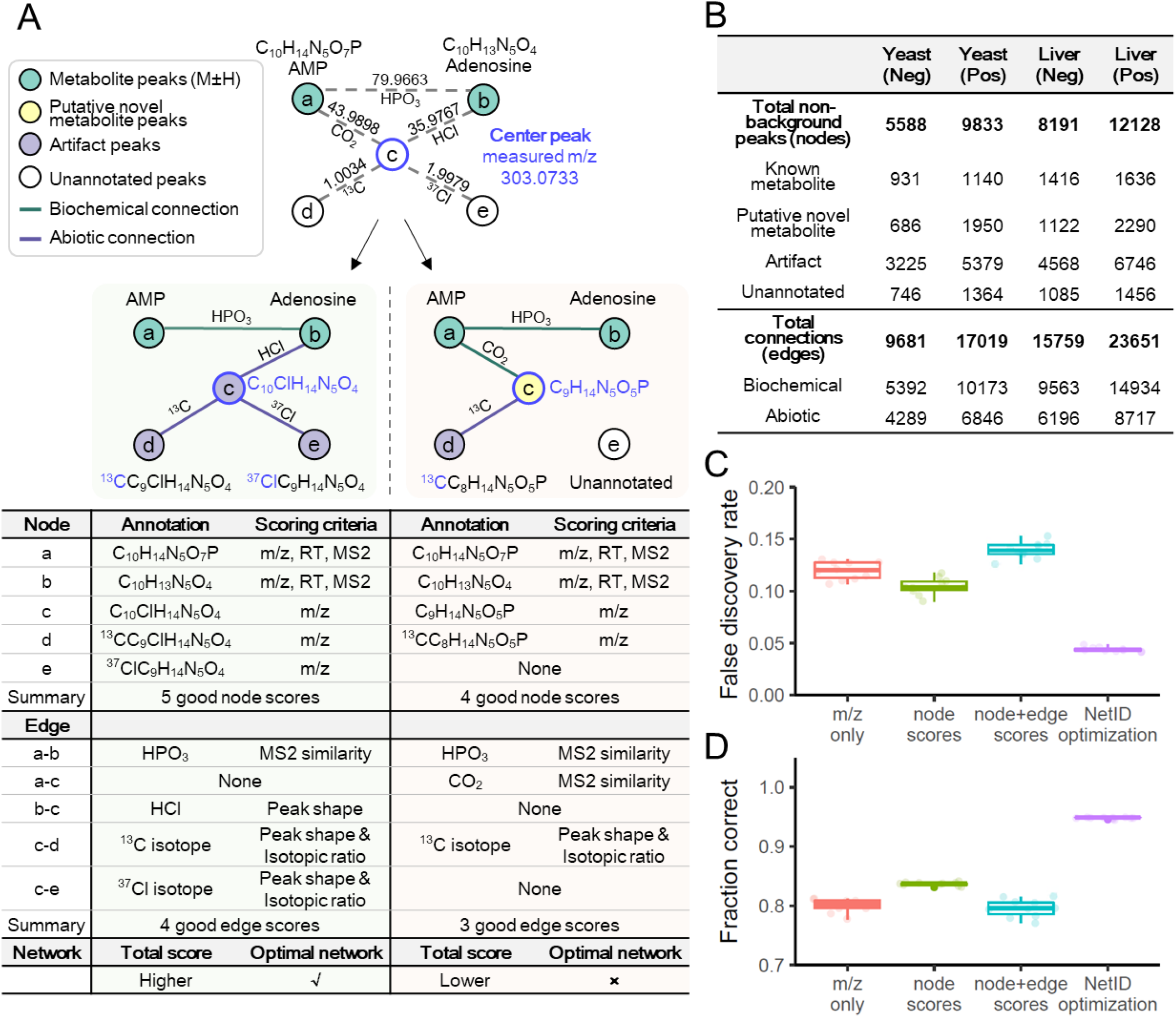
Utility of global network optimization. (A) An example network demonstrating the value of the global optimization step in NetID. Node *a* and node *b* match database formulae and are connected by an edge of phosphate (HPO3). Node *c* can be connected to either node *a* or node *b* through mutually incompatible annotations, resulting in two different candidate networks. The table below the two candidate networks shows the annotations and scoring criteria for each, with the left network preferred for more good node and edge annotations. (B) Summary table of NetID annotations of negative and positive mode LC-MS data from Baker’s yeast and mouse liver. (C) False discovery rate estimated using target-decoy strategy. Each individual data point (circle) is from a different randomized decoy library (n = 10 randomized libraries were tested). (D) Fraction of 314 manually curated “ground truth” annotations made correctly (using the 10 libraries). Boxes show median and interquartile range.

The left network has higher total node and edge annotation scores than the right network, and thus is selected by NetID. This selection makes sense to an experienced mass spectroscopist: the ^37^Cl isotope signature in node *e* indicates that node *c* should contain Cl. The power of NetID is that it automatically captures such logic, and uses the power of global computational optimization to extend such inferences across the network in an automated manner.

We applied the NetID algorithm to yeast and liver datasets, in both positive and negative ionization mode (Figure 2B, Supplementary Fig. 1A). Raw LC-MS data from replicate yeast or liver samples were analyzed together by peak-picking software (El-MAVEN^44^) to generate a single list of peaks consistently found for that sample type and ionization mode. Yeast data were MS1 only, while liver data included targeted MS2 spectra. Considering the example of negative mode yeast data with a total of 5,588 non-background peaks, in the candidate annotation step, roughly 1,600 potential formulae were assigned to 1,400 peaks, with about 200 peaks receiving multiple formula annotations. These nodes were connected by just over 50,000 potential edges. Edge extension expanded coverage to over 5,000 nodes with an average of twelve potential formulae each, highlighting the importance of scoring and network optimization to assign proper formulae. After scoring node and edge annotations, global network optimization settled on about 4,800 unique node annotations. About 20% of the annotated peaks were metabolites (formula corresponding to M±H monoisotopic peak existed in database), 14% were putative novel metabolites (formula not in database but with biochemical connection to a metabolite), and the rest were mass spectrometry phenomena, such as adducts, fragments, isotopes. Thus, after thorough background ion removal, we assign a few thousand peaks as likely metabolite ions and the majority as mass spectrometry artifacts. Orthogonal approaches such as credentialing via isotope labeling^27^ similarly assign the majority of peaks as mass spectrometry artifacts, but annotate fewer peaks as likely metabolites than NetID.

The roughly 5000 nodes were connected by about 10,000 edges. Two nodes share each edge, with each node connected by an average of four edges. These edges were roughly evenly split between biochemical and abiotic connections (Figure 2B, Supplementary Fig. 1A,B). More than 90% of annotated nodes fell into a single dominant connected network (Supplementary Fig. 1C), reflecting most peaks being connected to core metabolism. About 15% of peaks (737/5588), however, remained unannotated (Figure 2B). These unannotated peaks likely reflect deficiencies in our lists of allowed atom differences, including additional forms of mass spectrometry phenomena. For example, manual examination of the unconnected peaks revealed a dozen nickel adducts of known metabolites (Supplementary Table. 3). The annotated peaks included several hundred formulae for putative novel metabolites (Supplementary Fig. 2, Supplementary Data 1).

### Performance validation

We evaluated the performance of the NetID algorithm using the negative mode yeast data (MS1 only). We first employed a target-decoy estimation strategy, in which we intentionally introduce formulae with biologically unreasonable elements, and test whether our annotation strategy effectively avoids annotating peaks with these fake formulae^45,46^. Assessments were made using a number of different metabolite databases (HMDB^7^, YMDB^47^, PubChemLite.0.2.0^48^, and a subset of biopathway related entries in PubChemLite.0.2.0). As expected, the smaller databases yielded fewer false identifications. Importantly, across all of the databases, NetID more effectively selected appropriate formulae (lower false discovery rate) compared to methods considering m/z only, node scores only, or both node and edge scores but without global optimization (Figure 2C, Supplementary Fig. 3A).

As an orthogonal means of testing the algorithm, we manually curated 314 peaks as known annotations (Supplementary Data 1), and assessed the fraction annotated correctly. Across databases, NetID resulted in more accurate annotations of these gold standard peaks, with the number of incorrect annotations roughly an order of magnitude lower for NetID compared to node or combined node and edge scores without global optimization (Figure 2C, Supplementary Fig. 3B).

### Thiamine-derived metabolites

Among the putative novel metabolites in the yeast metabolomics dataset, we found three with total ion current > 10^5^ that are connected in a subnetwork around thiamine. Their formulae are C_12_H_16_N_4_O_2_S (thiamine+O), C_14_H_20_N_4_O_2_S (thiamine+C_2_H_2_O) and C_14_H_18_N_4_O_2_S, (thiamine+C_2_H_4_O) (Figure 3A, Supplementary Fig 4). These formula assignments and connections were initially obtained without MS2 spectra being available, reflecting the ability of NetID to make accurate formula assignments and connections based on MS1 data (combined with other peak attributes like retention time). While not found in HMDB, thiamine+O is documented in METLIN as a thiamine oxidation product, so we focused on the other two potential thiamine derivatives.

**Figure 3.**
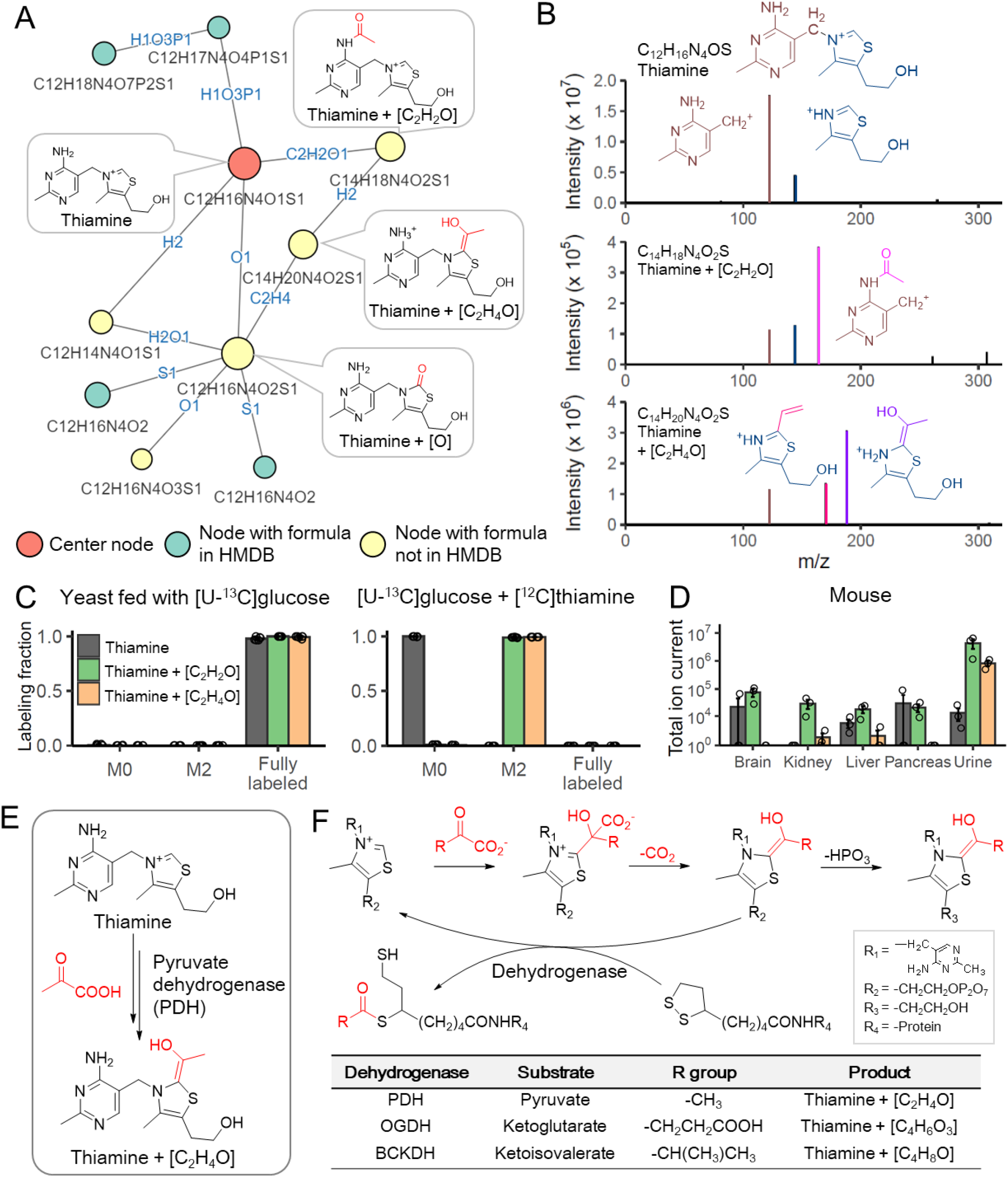
NetID reveals thiamine-derived metabolites in yeast. (A) Subnetwork surrounding thiamine. Nodes, connections, and formulae are direct output of NetID. Boxes with structures were manually added. (B) MS2 spectra of thiamine, thiamine+C_2_H_2_O, and thiamine+C_2_H_4_O, with proposed structures of the major fragments. (C) Labeling fraction of thiamine and its derivatives, in [U-^13^C]glucose with and without unlabeled thiamine in the medium (n = 5). (D) The thiamine derivatives are also found in mouse tissues and urine (n=3). (E) Proposed mechanism for formation of thiamine+C_2_H_4_O. Pyruvate dehydrogenase (PDH) decarboxylates pyruvate, and adds the resulting [C_2_H_4_O] unit (in red) to thiamine. (F) The same enzymatic mechanism occurs in oxoglutarate dehydrogenase (OGDH) and branched-chain α-ketoacid dehydrogenase complex (BCKDC), and generates thiamine+C_4_H_6_O_3_ and thiamine+C_4_H_8_O respectively. Bar represents mean values and error bar indicates s.d. in (C) and s.e. in (D).

MS2 spectra of the putative thiamine+C_2_H_2_O and thiamine+C_2_H_4_O contained characteristic thiamine fragments. Both contained a classical pyrimidine fragment, with thiamine+C_2_H_2_O also containing an acetylated pyrimidine fragment, leading to a probable structure (Figure 3A,B). The structural assignment is further supported by the presence of an unmodified thiazole fragment. In contrast, thiamine+C_2_H_4_O lacked a classical unmodified thiazole fragment, instead showing a thiazole+C_2_H_4_O fragment (and a fragment with further water loss) (Figure 3A,B).

Isotope tracing experiments further confirmed these two peaks contain thiamine. When fed [U-^13^C]glucose as sole carbon source, yeast synthesize thiamine *de novo*, resulting fully labeled thiamine species, with carbon counts matching the NetID formula assignments (Figure 3C). Adding unlabeled thiamine to the [U-^13^C]glucose culture media, yeast uptake the unlabeled thiamine, resulting in unlabeled thiamine and M+2 labeled thiamine+C_2_H_2_O and thiamine+C_2_H_4_O species. Although discovered in yeast, these are conserved metabolites, found also in mammalian samples (Figure 3D).

Acetylation is one of the 25 biochemical atom transformations allowed in NetID, while the addition of C_2_H_4_O is much less common biochemically. Accordingly, we looked into thiamine metabolism to explore how thiamine+C_2_H_4_O might be produced. Thiamine pyrophosphate is an important cofactor in pyruvate dehydrogenase (PDH, the entry step of carbohydrate to TCA cycle) (Figure 3E). The depyrophosphorylation product of thiamine pyrophosphate intermediate in PDH reaction yields thiamine+C_2_H_4_O (Figure 3F).

Based on this biochemical route, we realized that analogous products could be formed by α-ketoglutarate dehydrogenase (thiamine+C_4_H_6_O_3_) and branched-chain keto acid dehydrogenase (thiamine+C_4_H_8_O) (Figure 3F). Peaks at both of these exact masses were also experimentally observed, with isotope labeling results supporting their being thiamine-derived metabolites (Supplementary Fig. 5). Thus, NetID enabled the discovery of four novel thiamine-derived metabolites that were not present in metabolomics databases (Supplementary Table 4).

### N-glucosyl-taurine

We similarly carried out NetID annotation of a mouse liver dataset. We observed multiple putative novel metabolite peaks linked to taurine, by apparent glucosylation (+C_6_H_10_O_5_), palmitylation (+C_16_H_30_O) and transamination (+O-NH_3_) (Figure 4A, Supplementary Fig. 6). Like the thiamine-related peaks, these were initially correctly annotated without relying on MS2 data. While all were missing in HMDB, the latter two were found in METLIN: N-palmitoyl taurine (C_18_H_37_NO_4_S) and sulfoacetaldehyde (C_2_H_4_O_4_S). Pubchem contains an entry for N-glucosyl-taurine (C_8_H_17_NO_8_S) as a synthetic chemical but no database previously suggested it being a metabolite. To confirm the structure of the putative taurine glucosylation product (C_8_H_17_NO_8_S), we chemically synthesized N-glucosyl-taurine. Synthetic N-glucosyl-taurine matched the retention time and MS2 fragmentation pattern of the observed C_8_H_17_NO_8_S peak (Figure 4B,C). In liver samples of mice infused with [U-^13^C]glucose, C_8_H_17_NO_8_S appeared in M+6 form, suggesting active biosynthesis of N-glucosyl-taurine from circulating glucose (Figure 4D). N-glucosyl-taurine was not observed in yeast extract but was detected in multiple mouse tissues. Search for peaks matching the N-glucosyl-taurine MS2 spectra using MASST identified matches in both mouse and human samples^49^. Quantitation using the synthetic standard shows that liver has the highest level of glucosyl-taurine at ~170 μM (Figure 4E, Supplementary Fig. 7). This ranks among the few dozen most abundant liver metabolites.

**Figure 4.**
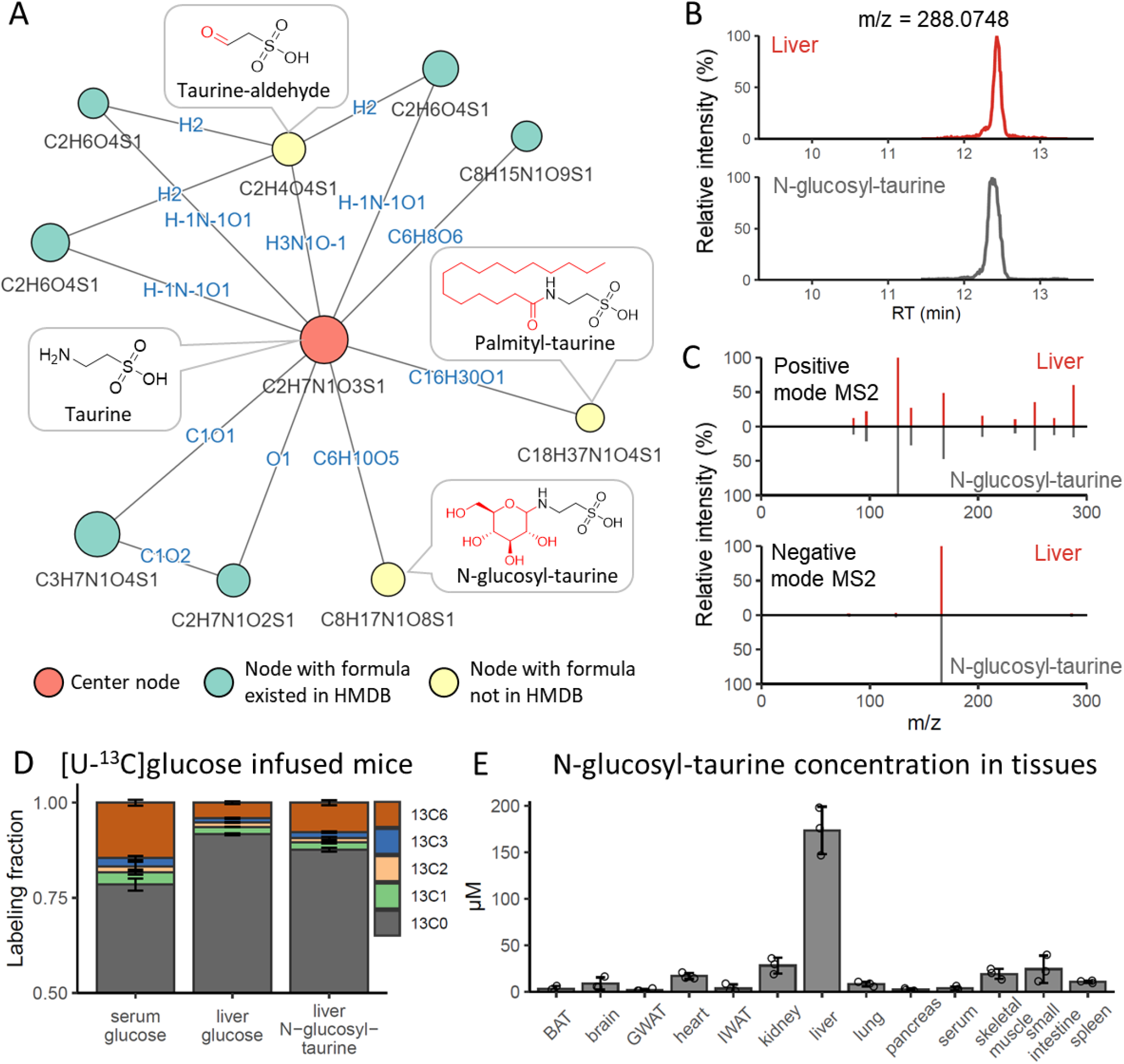
NetID discovers mammalian taurine derivatives. (A) Subnetwork surrounding taurine from mouse liver extract data. Nodes, connections, and formulae are direct output of NetID. Boxes with structures were manually added. (B) LC-MS chromatogram of N-glucosyl-taurine standard and the putative glucosyl-taurine from liver extract. (C) Top 10 abundant ion peaks in MS2 spectrum of glucosyl-taurine peak from liver extract (top), and synthetic N-glucosyl-taurine standard (bottom). (D) Isotope labeling pattern of putative glucosyl-taurine in mice, infused via jugular vein catheter for 2 h with [U-^13^C]glucose (n=3). (E) Absolute N-glucosyl-taurine concentration in murine serum and tissues (n=3). Bar represents mean values and error bar indicates s.d. in (D) and s.e. in (E).

## Discussion

The advent of LC-MS metabolomics revealed tens of thousands of metabolite peaks not matching known formulae, raising the possibility that the majority of metabolites remained to be discovered. While the biosphere likely contains many novel metabolites, it has been increasingly recognized that most peaks in typical untargeted metabolomics studies do not arise from novel metabolites, but rather mass spectrometry phenomena. The goal of comprehensively annotating untargeted metabolomics peaks with molecular formulae has, however, remained elusive.

One promising strategy for peak annotation involves building networks where nodes are LC-MS peaks (with associated molecular formulae) and edges are atom transformations linking the peaks. Here we advance this strategy by combining metabolomics knowledge with computational global optimization. We explicitly differentiate biochemical connections reflecting metabolic activity and abiotic connections arising from mass spectrometry phenomena. By formulating the peak annotation challenge as a linear program, we identify an optimal network in light of all observed peaks. Rather than weeding out peaks from mass spectrometry phenomena like adducts and natural isotopes, this approach takes advantage of the information embedded in them. It further provides traceable peak-peak relationships, which illuminate the basis for assigned formulae and suggest candidate structures.

Applying this approach to untargeted LC-MS data from yeast and liver samples, we assign formulae to roughly three-quarters of all non-background peaks. In each of positive and negative mode, the annotated peaks cover about 1000 known metabolites, with on average three mass peaks for every metabolite or putative metabolite (e.g. M+H plus two adduct or isotope peaks). This leaves a couple thousand unannotated peaks from each LC-MS run. Based on the observed ratio between peaks and metabolites, this likely correspond to hundreds (but not thousands) of unidentified metabolites. The number of actual novel metabolites being detected could potentially be more, due to isomers sharing the same formula as known metabolites, or less, due to novel adducts (e.g. nickel adducts, which we discovered via careful examination of the unannotated peaks) or other mass spectrometry phenomena. Importantly, this approach has already generated likely formulae for many hundreds of putative novel metabolites (Supplementary Fig. 2, Supplementary Data 1), including five species for which we assign structures (Figure 3, 4).

A key benefit of molecular network-based annotation is the ability to assimilate steadily new information^28,29^. MS2 data, which can be collected in either targeted or data-dependent mode, provide fragmentation information to facilitate peak annotation. Due to the global network optimization approach, new MS2 spectra facilitate not only annotation of the ions being fragmented, but also other peaks throughout the network. Other data types can be seamlessly added. For example, compound class categorization based on MS2 data^50^ or retention time prediction^51,52^ can be added to score nodes. Isotope labeling similarity upon feeding different “heavy” nutrients could potentially be added to score edges. Global optimization integrates new information comprehensively with prior knowledge to arrive at optimal annotations. Cycles of experimentation and machine learning hold the potential to identify most unknown metabolites over the coming decade, eventually providing a robust blueprint of the metabolome (Figure 5).

**Figure 5.**
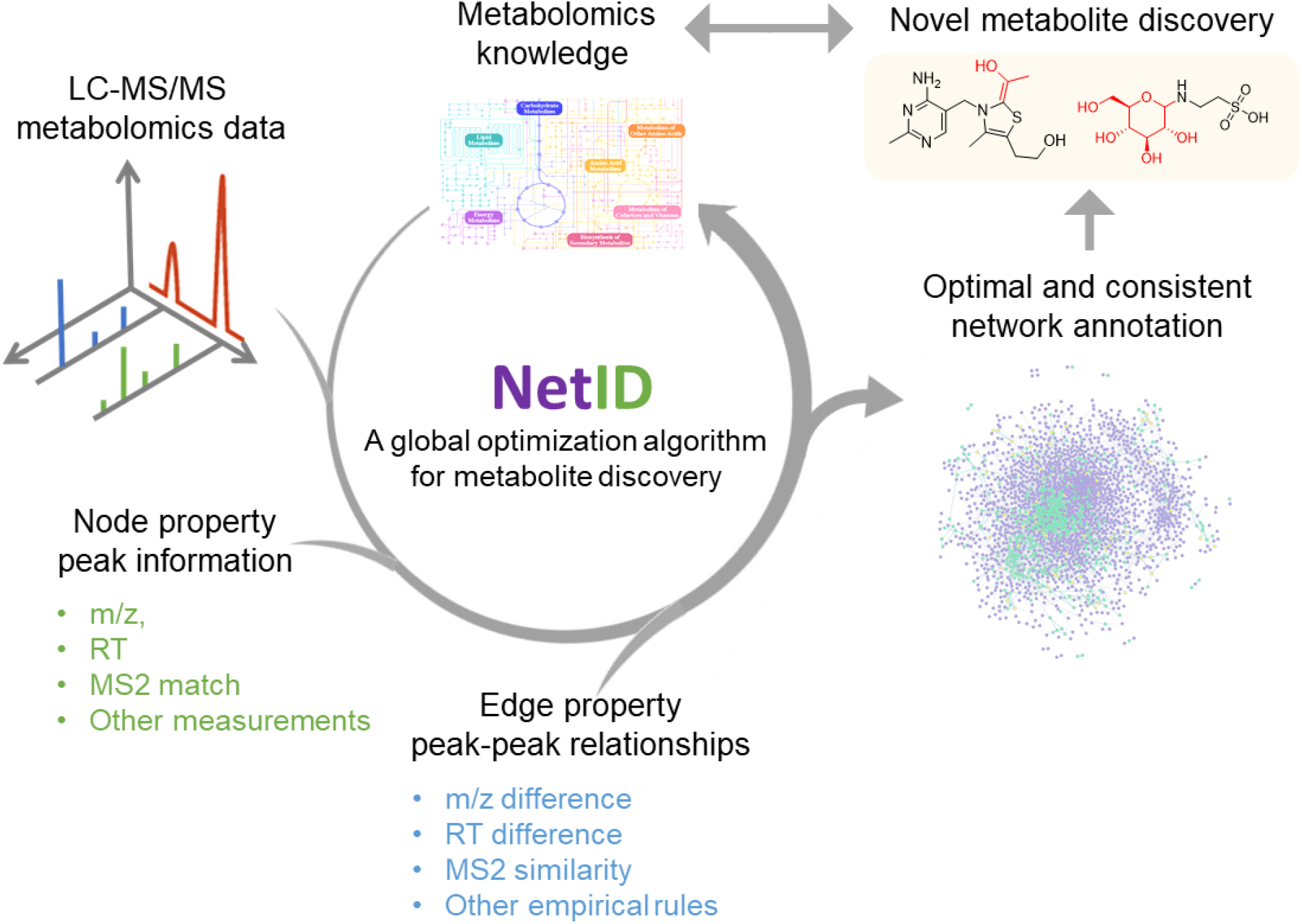
NetID applies global optimization for metabolomics data annotation and metabolite discovery.

## Methods

### Yeast metabolomics sample preparation and isotope labeling

*S. cerevisiae* strain FY4 was grown for at least 10 generations in minimal essential media containing 0.4% [U-^12^C] or [U-^13^C]glucose and 10 mM ammonium sulfate with or without 0.4 mg/L thiamine hydrochloride^53^. Then, in mid-exponential phase, 5 mL culture broth (OD_600_ = 0.80) was filtered and metabolites were extracted using 1 mL extraction buffer (40:40:20:0.5 acetonitrile:methanol:water:formic acid), followed by adding 88 μL neutralization buffer (15% NH_4_HCO_3_). The extracts were kept at −20℃ for at least 15 min to precipitate protein before centrifuging at 16,000 g for 10 min. The supernatant was used for LC–MS analysis.

### Murine metabolomics sample preparation and intravenous infusion experiment

Animal studies followed protocols approved by the Princeton University Institutional Animal Care and Use Committee. Twelve-month-old female wild-type C57BL/6 mice (The Jackson Laboratory, Bar Harbor, ME) on standard mouse chow diet were sacrificed by cervical dislocation and tissues quickly dissected and snap frozen in liquid nitrogen with precooled Wollenberger clamp. Frozen samples from liquid nitrogen were then transferred to −80°C freezer for storage. To extract metabolites, frozen liver tissue samples were first weighed (~ 20 mg each) and transferred to 2 mL round-bottom Eppendorf Safe-Lock tubes on dry ice. Samples were then ground into powder with a cryomill machine (Retsch, Newtown, PA) for 30 seconds at 25 Hz, and maintained at cold temperature using liquid nitrogen. For every 25 mg tissues, 922 uL extraction buffer (as above) was added to the tube, vortexed for 10 seconds, and allowed to sit on ice for 10 minutes. Then 78 μL neutralization buffer was added and the samples vortexed. The samples were allowed to sit on ice for 20 minutes and then centrifuged at 16,000 g for 25 min at 4°C. The supernatants were transferred to another Eppendorf tube and centrifuged at 16,000 g for another 25 min at 4°C. The supernatants were transferred to glass vials for LC-MS analysis. A procedure blank was generated identically without tissue, which was used later to remove the background ions.

Detailed methods for intravenous infusion of mice have been described previously^54^. Briefly, *in vivo* infusions were performed on 12–14-week-old C57BL/6 mice pre-catheterized in the right jugular vein (Charles River Laboratories). Mice were kept fasted for 6 h and then infused for 2.5 h with [U-^13^C]glucose (200 mM, 0.1 μL/min/g). The mouse infusion setup (Instech Laboratories) included a tether and swivel system so that the animal had free movement in the cage. Venous samples were taken from tail bleeds. At the end of the infusion, the mouse was euthanized by cervical dislocation and tissues were collected and extracted as above. Serum metabolites were extracted by adding 100 μl methanol to 5 μL of serum and centrifuging for 20 min. The supernatant was used for LC–MS analysis.

### Glucosyl-taurine synthesis

Glucosyl-taurine synthesis was carried out following previous literature reports with slight modifications^55^. In brief, dry methanol was obtained by distillation of HPLC-grade methanol (Fisher; HPLC grade 0.2 micron filtered) over CaH_2_ (Acros Organics; ca. 93% extra pure, 0-2 mm grain size). A flame-dried round-bottom flask equipped with a reflux condenser and stir bar was charged with 2.0 g taurine (Alfa Aesar; 99%), 3.1 g D-glucose (Acros Organics; ACS reagent), and 80 mL of dry methanol. This mixture was sonicated under an inert atmosphere for 30 min before being returned to the manifold for the reaction. To the fine-suspension of taurine and glucose in dry methanol at room temperature, 4.0 mL 5.4 M sodium methoxide in methanol (Acros Organics) was added via glass syringe. At this point, the suspension began to dissolve and after 30 minutes, gave a clear and colorless solution. The solution was stirred vigorously under an inert atmosphere for 72 hours, which resulted in a faint peach-colored solution. This solution was chilled to 0 °C, and ~200 mL of absolute ethanol (200 proof) was added and precipitation was allowed to occur at this temperature for 30 minutes. Solvent was then removed by filtration over a glass filter (medium porosity), and washed with ~100 mL of absolute ethanol, affording a fine pale-yellow powder (2.4 g; crude material).

NMR was carried out to validate the structure of synthesized N-glucosyl-taurine. Selective TOCSY experiments using DIPSI2 spin-lock and with added chemical shift filter^56^ were run on a Bruker Avance III HD NMR spectrometer equipped with a custom-made QCI-F cryoprobe (Bruker, Billerica, MA) at 800 MHz and at 295 K controlled temperature. The sample was dissolved in DMSO-d6. The spectra shown on the plots are results of 200 ms SL mixing, 8 scans each. Data processing (MNova v.14, Mestrelab Research S.L., Santiago de Compostela, Spain) included zero filling, 1 Hz Gaussian apodization, phase- and baseline correction. NMR analysis suggests that the final crude material contains 5.2% N-glucosyl-taurine mixed with unreacted substrates (Supplementary Fig. 8).

### LC-MS and LC-MS/MS

LC separation was achieved using a Vanquish UHPLC system (Thermo Fisher Scientific) with an Xbridge BEH Amide column (150×2mm, 2.5 μm particle size; Waters). Solvent A is 95:5 water: acetonitrile with 20 mM ammonium acetate and 20 mM ammonium hydroxide at pH 9.4, and solvent B is acetonitrile. The gradient is 0 min, 90% B; 2 min, 90% B; 3 min, 75%; 7 min, 75% B; 8 min, 70%, 9 min, 70% B; 10 min, 50% B; 12 min, 50% B; 13 min, 25% B; 14 min, 25% B; 16 min, 0% B, 20.5 min, 0% B; 21 min, 90% B; 25 min, 90% B. Total running time is 25 min at a flow rate of 150 μl/min. LC-MS data were collected on a Q-Exactive Plus mass spectrometer (Thermo Fisher) operating in full scan mode with a MS1 scan range of m/z 70-1000, and resolving power of 160,000 at m/z 200. Other MS parameters are as follows: sheath gas flow rate, 28 (arbitrary units); aux gas flow rate, 10 (arbitrary units); sweep gas flow rate, 1 (arbitrary units); spray voltage, 3.3 kV; capillary temperature, 320°C; S-lens RF level, 65; AGC target, 3E6 and maximum injection time, 500 ms.

MS2 spectra were collected in targeted mode using the PRM function at 25 eV HCD energy with other instrument settings being resolution 17500, AGC target 10^6^, Maximum IT 250 ms, isolation window 1.5 m/z. Targeted MS2 data for the thiamine related metabolites were collected for structural identification using similar parameters as above except the HCD energy were set at 20, 35, and 50 eV in a step-CE mode.

### Data prepossessing

LC-MS raw data files (.raw) were converted to mzXML format using ProteoWizard^57^. El-MAVEN (version 7.0) was used to generate a peak table containing m/z, retention time, and intensity for peaks. Parameters for peak picking were the defaults except for the following: mass domain resolution is 10 ppm; time domain resolution is 15 scans; minimum intensity is 1000; minimum peak width is 5 scans. The resulting peak table was exported to a .csv file. Redundant peak entries due to imperfect peak picking process are removed if two peaks are within 0.1 min and their m/z difference are within 2 ppm. Background peaks are removed if its intensity in procedure blank sample is > 0.5-fold of that in biological sample.

Targeted MS2 data were extracted from the mzXML files using lab-developed Matlab code (Supplementary Note 1). MS2 spectra may contain interfering product ions from co-eluting isobaric parent ions. These interfering product ions were removed by examining the extracted ion chromatogram (EIC) similarity between the product ions in MS2 data and the parent ion in MS1 data. A Pearson correlation coefficient of 0.8 was used as a cutoff to retain those product ions that has similar EIC as the parent ion. The cleaned MS2 data were exported to Excel files for data input. Although the provided workflow uses targeted MS2 data as input, NetID as currently configured can also handle data-dependent MS2 data, but additional parsing software (under development) is need to convert the large primary data-dependent MS2 files into the NetID input format.

### Data input

NetID algorithm requires three types of input files: a peak table (in .csv format) recording m/z, retention time, and intensity for peaks; an atom difference rule table (in .csv format), which in our case consisted of a list of 25 biochemical atom differences and 59 abiotic atom differences which together define all connections in the network (Supplementary Table 1, 2), and metabolite information files containing structures, formulae, m/z and MS2 spectra of metabolites from the selected metabolomics database (e.g. HMDB) including retention times of known metabolites under the relevant LC conditions. MS2 data input is optional. For the analysis of yeast and mouse liver datasets in Figure 2–4, structures, formulae, m/z and MS2 spectra of metabolites were obtained from the Human Metabolome Database (HMDB, version 4.0) and retention times of selected metabolites were determined through running authentic standards using the above-mentioned LC-MS method (Supplementary data 2). For yeast, no MS2 were used in NetID analysis (MS2 were used post hoc to confirm certain annotations). For liver, targeted MS2 spectra were used (1479 positive and 803 negative ionization mode spectra experimentally obtained for previously identified peaks of > 10^5^ intensity^58^).

### Candidate node and edge annotations

The first module of NetID algorithm is to make candidate annotations for seed nodes, assign candidate annotations for other nodes, and assign candidate edges in the network. Each peak is a node in the network. We compare the experimentally measured m/z for each node to those of all metabolite formulae in the selected metabolomics database (e.g. HMDB). When the m/z difference is within a predefined tolerance (e.g. 10 ppm), candidate formulae and IDs are assigned to the node, and this node is defined as a primary seed node. Note that assignments to seeds are candidate annotations. A primary seed node can contain multiple candidate formulae and IDs if all are within the m/z difference range.

Edges connect two nodes via gain or loss of specific atoms. We provided a list of 25 biochemical atom differences and 59 abiotic atom differences which together define all connections in the network (Supplementary Table 1, 2, Supplementary Data 2). Let each of these differences be denoted by *D*_*i*_. For each node *u*, if there is a node *v* such that the difference in the measured m/z of the nodes matches one of the those in the list of atom mass differences within a predefined *m*/*z*_*tol*_ (e.g. 10 ppm), we add an edge between *u* and *v*. That is, if *u*_m/z_ and *v*_m/z_ are the experimentally measured m/z for the peaks corresponding to nodes *u* and *v* respectively (assuming *v*_m/z_ > *u*_m/z_ for simplicity), then there is an edge between these nodes if there is some difference *D*_*i*_ such that

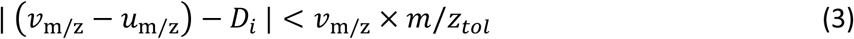

If *D*_*i*_ is an abiotic difference, in order to add an edge, it is additionally required that the retention time between two nodes should be within a predefined *RT*_*tol*_ (e.g. 0.2 min). That is, if *u*_RT_ and *v*_RT_ are the retention times for *u* and *v* respectively, then it is required that

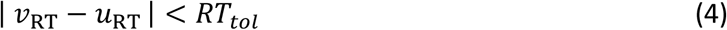

For each node, its candidate formulae set will expand due to extending formulae from its neighboring nodes through the edge atom differences. For example, when applying the atom difference of edge (*u*, *v*) on the formula assigned to primary seed node *u*, we can derive a new candidate formula for the connected node *v*. If the derived formula’s calculated m/z is within a predefined m/z tolerance (e.g. 5 ppm) of node *v*’s measured m/z, then a new candidate formula is added for node *v*. Iterating the process to all candidate formulae of node *u* through edge (*u*, *v*) will further expand candidate formulae for node *v*.

We apply the above extension process to formulae of all primary seed nodes through atom difference edges, and these new candidate formulae themselves can be used for another round of extension. Note that a primary seed node will be treated as the rest of nodes during the subsequent rounds of extension, and may as well be assigned with new formulae. To avoid duplicated efforts in the extension process, we allow formulae of primary seed nodes and biotransformed formulae thereof to be extended through both biotransformation and abiotic atom difference edges, and do not allow abiotic candidate formulae be further extended through biotransformation atom difference edges. The default extension process includes two rounds of biotransformation edge extensions and three rounds of abiotic edge extensions.

Each candidate node annotation is defined as (i) metabolite, (ii) putative novel metabolite, or (iii) artifact (nodes can also be unannotated). Specifically, if the elemental formula corresponding to the (de)protonated ion of a monoisotopic peak is found in the employed metabolomics database, this node is defined as a metabolite. If the formula is not found at the employed database, but the node is connected to a metabolite via biochemical connection(s), the node is defined as a putative novel metabolite. Finally, if the node connected only via abiotic connections such as adduct, fragment, or isotope connection(s), it is defined as an artifact. As currently configured, NetID defines metabolite peaks exclusively as (de)protonated ions. In the case that a (de)protonated ion peak is not detected, but related adducts are (e.g. [M+Na]^+^), the adducts will remain unannotated (or be misannotated), as there is no procedure for annotating adducts lacking (de)protonated ion peaks.

### Scoring in NetID

The second module of NetID algorithm is to score every candidate node and edge annotation assigned in the candidate annotation step.

The node scoring system aims to assign high scores to annotations that align observed ion peaks with known metabolites based on m/z, retention time, MS2, and/or isotope abundances. Let the set of candidate annotation for node *u* be denoted as {*a*_1_ … *a*_*i*_ … *a*_*m*_}. For each node *u* and each of its candidate annotation *a*_*i*_ , let S(*u*, *a*_*i*_) denotes the score of candidate annotation *a*_*i*_ for node *u*. S(*u*, *a*_*i*_) is the sum of seven different scoring components, including (a) S_m/z_, a negative score evaluating the difference of between measured m/z and the calculated m/z of assigned molecular formula; (b) S_RT_, a positive score if the measured RT for the peak corresponding to node *u* matches to a known standard; (c) S_MS2_, a positive score if the measured MS2 spectrum of node *u* matches the database MS2 spectrum of annotation *a*_*i*_; (d) S_database_, a positive score if the annotated formula *a*_*i*_ exists in the employed metabolomics database; (e) S_missing_isotope_, a negative score if an expected isotopic peak is missing; (f) S_rule_, a negative score if annotation *a*_*i*_ violates basic chemical rules; (g) S_derivative_, a positive score if the annotation *a*_*i*_ is derived from a parent peak with a high score annotation. For details, see Supplementary Note 2.

The edge scoring system aims to assign high scores to edge annotations that correctly capture biochemical connections between metabolites (based on MS2 spectra similarity) and abiotic connections between metabolites and their mass spectrometry phenomena derivatives. Biochemical, isotope, adduct, and neural loss edge annotations are the most common types. We also score other less common abiotic connection types appeared in orbitrap data, including oligomers, multi-charge species, heterodimers, in-source fragments of known or unknown metabolites^59^, and ringing artifact peaks surrounding high intensity ions^26,60^.

Suppose we consider two nodes *u* and *v* that are connected by an edge (*u*, *v*). For each pair of nodes *u* and *v* such that there is an edge (*u*, *v*), let the set of candidate formula for node *u* and *v* be denoted as {*a*_1_ … *a*_*i*_ … *a*_*m*_} and {*b*_1_ … *b*_*j*_ … *b*_*n*_}, respectively, and let the set of candidate atom differences for edge (*u*, *v*) be {*D*_1_ … *D*_*k*_ … *D*_*l*_}. Let S(*u*, *ν*, *a*_*i*_, *b*_*j*_, *D*_*k*_) be the score of choosing candidate formula *a*_*i*_ for node *u*, candidate formula *b*_*j*_ for node *v* and candidate atom difference *D*_*k*_ for edge (*u*, *v*). Note that S(*u*, *ν*, *a*_*i*_, *b*_*j*_, *D*_*k*_) is set to be 0 if atom difference *D*_*k*_ does not represent the formula difference of *a*_*i*_ and *b*_*j*_.

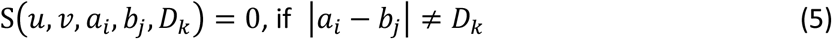

S(*u*, *ν*, *a*_*i*_, *b*_*j*_, *D*_*k*_) is the sum of four different scoring components, including (a) S_MS2_similarity_, a positive score defined for biochemical edges if node *u* and *v* have experimental measured MS2 spectra, and they share MS2 similarity; (b) S_co_elution_, a negative score defined for abiotic edges, if the RT of two connected nodes differ more than a threshold (e.g. 0.05 min); (c) S_type_(*u*, *ν*, *a*_*i*_, *b*_*j*_, *D*_*k*_), a non-negative score defined for all edges, depending on the connection type of edge, which is defined by *D*_*k*_, including biotransformation, adduct, isotope, and fragment edges, and optionally including oligomer and multi-charge species, heterodimer, in-source fragments and ringing artifacts edges; (d) S_*isotope*_*intensity*_, a negative score defined for isotopic edges (a type of abiotic edge) if the measured isotope peaks deviate from expected natural abundance. For details, see Supplementary Note 2.

### Global network optimization using linear programing

The third module of NetID algorithm is to perform global network optimization. Using scores assigned for each candidate node and edge annotation, our goal is to find formula annotations for each node and edge so as to maximize the sum of their scores across the network under the constraints that each node is assigned a single annotation, and that the network annotation is consistent. When a node has multiple candidate node annotations shared with same formula (e.g. isomers), the one with highest score (better MS2 match or RT match) is selected. When equal scores happen, the candidate annotation that appears first in the metabolite list from database is reported as a default. We use linear programming to solve this optimization problem, as described next.

For each node *u* and each of its candidate formula *a*_*i*_, we define a node binary decision variable 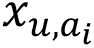 to denote whether candidate formula *a*_*i*_ is selected as the annotation for node *u*. That is,

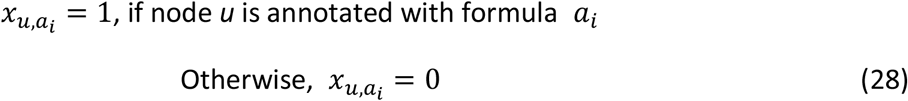

For each edge, we define a binary decision variable 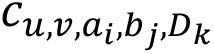 to denote whether candidate formulae *a*_*i*_ and *b*_*j*_ are chosen for nodes *u* and *v* , and the candidate atom difference *D*_*k*_ corresponds to the formula difference of candidate formulae *a*_*i*_ and *b*_*j*_ of the connected nodes *u* and *v*. That is,

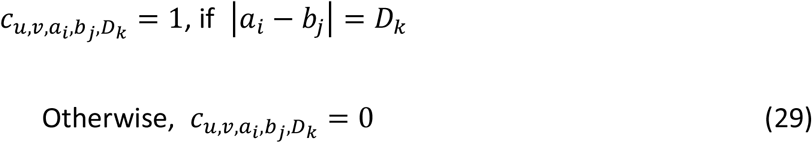

We constrain the optimization so that each node has a single annotation, and an edge exists only if the atom difference of that edge annotation matches the formula difference of nodes. For computational purposes, nodes may also receive “blank” or “no formula” annotation (we refer to such nodes in elsewhere in the text as “unannotated”). The node and edge binary variables must satisfy

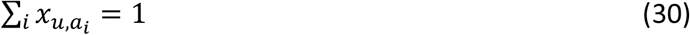

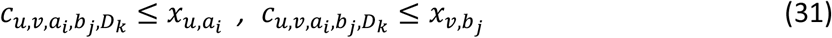

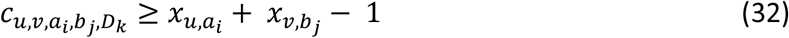

For all variables defined above, we add the constraints that they are either 1 or 0, representing the candidate annotation is selected or not selected, respectively, in the network.

With scores of each candidate node and edge annotation, the objective for the optimization is to determine all variables *x*_*u*,*a*_ and *C*_*u*,*ν*,*a*,*b*,*D*_ so as to maximize the sum of all node scores and edge scores in a network while satisfying the above constraints.

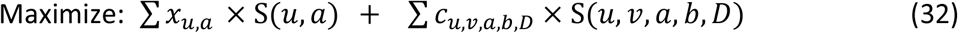

The optimization result provides a string of binary numbers that denote if a candidate node or edge annotation is selected for the global optimal network. IBM ILOG CPLEX Optimization Studio (Version 12.8.0 or later) is used to solve the linear programing problem. For the reported datasets and using the default parameter settings, optimization finishes within an hour on a standard laptop (Supplementary table 5). Depending on the number of peaks in data tables, the entries in the atom difference tables, the choice of reference compound databases, and the parameters involved in scoring, runtimes during internal testing ranged from minutes to hours.

### Evaluation of NetID

After running the candidate annotation and network extension process as usual for NetID, we compared four different annotation selection methods: (i) m/z only, selecting the candidate annotation with closest m/z to the measured m/z; (ii) node scores, selecting the candidate annotation with highest candidate node score (as per usual NetID scoring rules); (iii) node + edge scores, adding half of a candidate edge score to each node score of the two connected candidate node annotations, and selecting the candidate annotation with highest combined score; (iv) NetID optimization, using the candidate node and edge score, and selecting the candidate annotation from global optimization.

We employed a target-decoy strategy to estimate false discovery rate^45,46^. The target library is the compound library we use for annotation, including HMDB (human metabolomics database), PBCM (PubChemLite.0.2.0)^48^, PBCM_BIO (a subset of biopathway related entries in PubChemLite.0.2.0) and YMDB (yeast metabolomics database)^47^. The decoy formula was generated by adding an implausible element adduct to a formula from target library. These implausible elements are those not in any formulae in database, namely, He, Be, Ne, Sc, Kr, Rb, Sr, Y, Zr, Nb, Mo, Ru, Rh, Pd, Ag, Cd, In, Sn, Sb, Te, Xe, Cs, Ba, La, Ce, Pr, Nd, Sm, Eu, Gd, Tb, Ho, Er, Tm, Yb, Lu, Hf, Ta, W, Re, Os, Ir, Pt, Au, Hg, Tl, Pb, Bi, Th and U. Applying the decoy formula generation process (i.e. adding a single randomly selected implausible element in place of hydrogen) to all unique formulae in a target compound library gives the corresponding decoy library. The combined target library and decoy library were used for annotation. Any annotation containing an implausible element is considered a false positive (FP) result. The number of FP results from target library is estimated to be similar to that from decoy library, because 1:1 ratio of target formulae and decoy formulae were used. That is FP_target_library_ ≈ FP_decoy_library_. Using the combined target-decoy library, the false discovery rate (FDR) is estimated to be FP_decoy_library_ / (T - FP_decoy_library_), where T is the total number of annotations. The decoy library generation process was repeated ten times for each database.

We manually annotated 314 peaks in the yeast negative mode dataset (Supplementary data 1). Using these annotations as a ground truth, we evaluated the fraction of correct annotation for the four different annotation methods above. Peak annotations matching to the ground truth formulae, including adduct, isotope formulae, are counted as correct, and peaks that are not annotated or their annotations did not match are counted as incorrect. The annotation process used 1:1 target-decoy library and was repeated 10 times as above.

### Visualization

We provide an interactive Shiny R app to visualize and explore the NetID output network. In addition, NetID outputs two .csv files (cyto_node.csv and cyto_edges.csv) that are compatible with the general network visualization software Cytoscape. The interactive Shiny R app and detailed user guide are available at GitHub (https://github.com/LiChenPU/NetID).

### Data availability

All raw data, including the yeast, the mouse studies, the ^13^C studies and over 2000 targeted MS2 files collected from the liver data in mzXML formats were deposited in MassIVE (ID = MSV000087434). MS2 spectra of reported novel metabolites are included in Supplementary Data 3. R code for generating NetID statistics and performing FDR analysis in Figure 2 and Supplementary Fig 1 are provided in GitHub (https://github.com/LiChenPU/NetID). Peak table for yeast data negative mode, including NetID annotation results, putative novel metabolite list, and manual curation results are provided in Supplementary Data 1. Exemplary peak table from the yeast dataset, atom difference rule table, HMDB compound list (customized to contain relevant information), and an in-house retention time list for known metabolites are provided in Supplementary Data 2.

### Code availability

NetID was developed mainly in R, and used a mixture of IBM ILOG CPLEX Optimization Studio, Matlab and Python. NetID code and example files are available for non-commercial use in GitHub at https://github.com/LiChenPU/NetID, under the GNU General Public License v3.0. User guide, parameter explanation and pseudocode are provided in Supplementary Notes 1, 2, and 3.

## Acknowledgement

This work was supported by a Department of Energy (DOE) grant (no. DE-SC0012461 to J.D.R.), the Center for Advanced Bioenergy and Bioproducts Innovation (grant no. DE-SC0018420, subcontract to J.D.R.) and NIH grant R50CA211437 to W.L. M.R.M is funded by the Howard Hughes Medical Institute and Burroughs Wellcome Fund via the PDEP and Hanna H. Gray Fellows Programs. We thank Istvan Pelczer at NMR facility of Department of Chemistry at Princeton University for the NMR analysis, the Metabolomics & Lipidomics Mass Spectrometry Core Facility of IMIB at Fudan University for additional mass spectrometry support, and X. Su for scientific discussion and help. The Center for Advanced Bioenergy and Bioproducts Innovation and the Center for Bioenergy Innovation are both U.S. Department of Energy Bioenergy Research Centers supported by the Office of Biological and Environmental Research in the DOE Office of Science. Any opinions, findings, and conclusions or recommendations expressed in this publication are those of the authors and do not necessarily reflect the views of the U.S. Department of Energy.

## Author contributions

L.C., M.S. and J.D.R. conceived the project. L.C., X.X. and Z.C. wrote the NetID algorithm code. W.L., L.W., X.Z., A.C. M.M. performed experiments on mouse. L.W., W.L. and L.C. performed experiments on yeast. L.C., W.L., L.W. and X.X. analyzed LC-MS and LC-MS/MS data. X.T., A.M. and Y.S. contributed to coding development. B.K., A.M.L., and S.R.C. synthesized taurine-related compounds. L.C. and J.D.R. wrote the manuscript. All authors discussed the results and commented on the manuscript.

