## Supplementary for "Metabolite discovery through global annotation of untargeted metabolomics data"

#### Supplementary information

##### Supplementary Figures:

**Supplementary Figure 1.** Characterization of NetID network
**Supplementary Figure 2.** Examples of putative metabolites in yeast negative-mode dataset
**Supplementary Figure 3.** Evaluation of annotation false discovery rate (FDR) and fraction gold-standard peaks annotated correctly using different reference databases
**Supplementary Figure 4.** Subnetwork surrounding thiamine with additional known structures
**Supplementary Figure 5.** Evidence for the additional thiamine-derived metabolites
**Supplementary Figure 6.** Subnetwork surrounding taurine with additional known structures
**Supplementary Figure 7.** Glucosyl-taurine is a liver metabolite, not *ex vivo* reaction product
**Supplementary Figure 8.** SelTOCSY NMR confirmation of the structure of the chemically synthesized N-glucosyl-taurine

##### Supplementary Tables:

**Supplementary Table 1.** List of biochemical atom differences
**Supplementary Table 2.** List of abiotic atom differences
**Supplementary Table 3.** Examples of nickel-related peaks
**Supplementary Table 4.** Search results of reported novel metabolites in compound databases
**Supplementary Table 5.** Memory and run-time used in NetID

##### Supplementary Notes:

**Supplementary Note 1.** NetID user guide
**Supplementary Note 2.** NetID scoring
**Supplementary Note 3.** NetID pseudocode

##### Supplementary Data (in separate excel files):

**Supplementary Data 1.** NetID annotation for the yeast negative-mode dataset
**Supplementary Data 2.** Exemplary peak table, atom difference rule table and HMDB metabolite information
**Supplementary Data 3.** MS2 spectra of measured novel metabolites

A

| | | Yeast (Neg) | Yeast (Pos) | Liver (Neg) | Liver (Pos) |
|---|---|---|---|---|---|
| **Total non-background peaks** | | 5588 | 9833 | 8191 | 12128 |
| **Seed annotation** | **Seed nodes** | 2000 | 3092 | 2957 | 3746 |
| | with single candidate formula | 1731 | 2677 | 2589 | 3356 |
| | with multiple candidate formula | 269 | 415 | 368 | 390 |
| | **Total candidate formulas for seed nodes** | 2309 | 3554 | 3377 | 4176 |
| | **Candidate edges** | 57877 | 142377 | 106228 | 192141 |
| | Biochemical | 37075 | 96769 | 66011 | 114013 |
| | Abiotic | 20802 | 45608 | 40217 | 78128 |
| **Propagation** | **Nodes with candidate formula annotations** | 5253 | 9393 | 7701 | 11749 |
| | **Nodes without any annotations** | 335 | 440 | 490 | 379 |
| | **Total candidate formula annotations** | 61639 | 157774 | 98608 | 176958 |
| | Metabolite | 2292 | 3548 | 3346 | 4169 |
| | Putative metabolite | 11598 | 28868 | 19681 | 31515 |
| | Artifact | 47749 | 125358 | 75581 | 141274 |

B

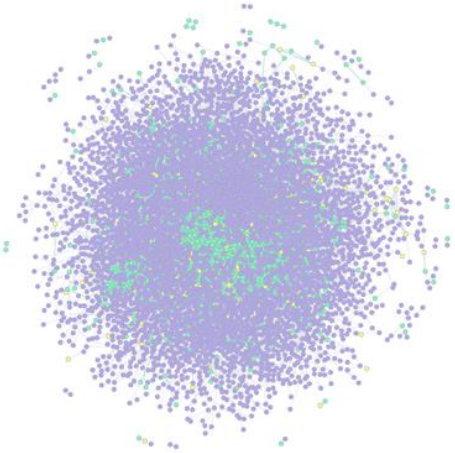

C

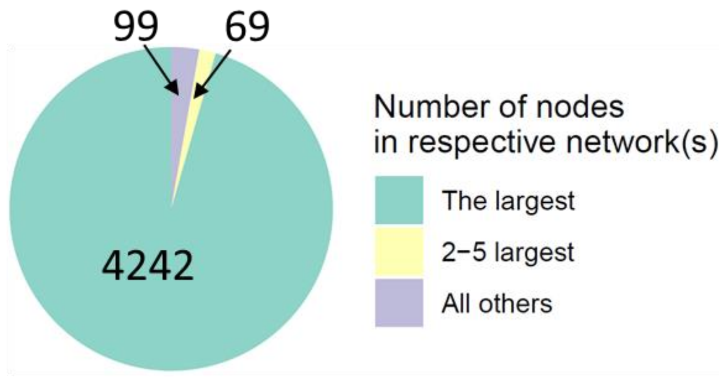


**Supplementary Figure 1.** Characterization of NetID network. (A) Summary table of the candidate annotation step in NetID workflow. (B) Visualization of the optimal network obtained from negative mode LC-MS analysis of Baker's yeast, containing 4851 nodes and 9699 connections. Metabolite and putative metabolite peaks are in green and artifact peaks in purple. (C) Connectivity of NetID network from the yeast negative-mode dataset.

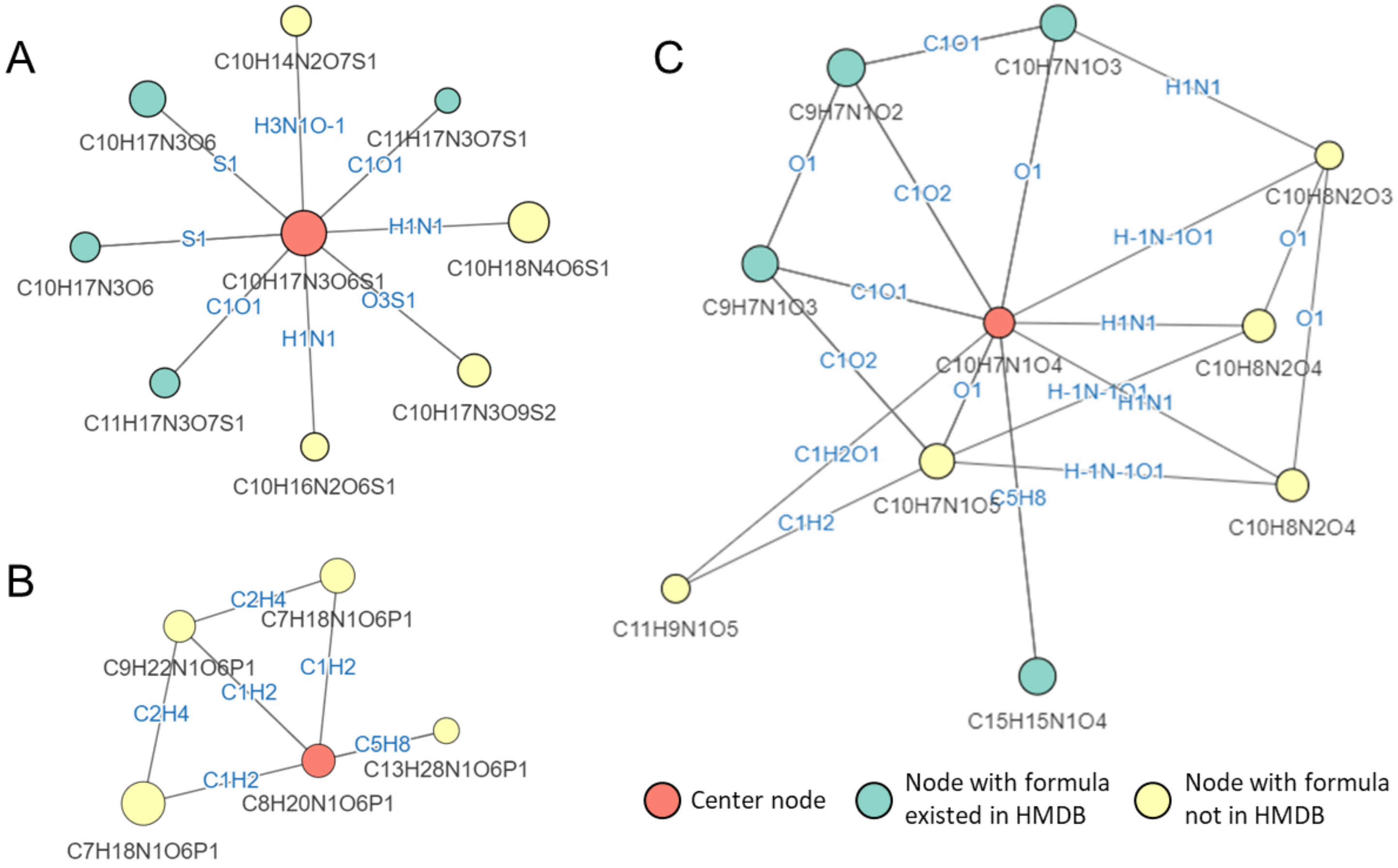


D

| panel | peak_id | medMz | medRt | log10_inten | formula | Δppm | annotation |
|---|---|---|---|---|---|---|---|
| A | 230 | 322.0946 | 13.68 | 6.49 | C10H18N4O6S1 | 0.33 | C10H17N3O6S1 + H1N1 -> C10H18N4O6S1 |
| | 1517 | 387.0407 | 13.52 | 5.24 | C10H17N3O9S2 | -0.21 | C10H17N3O6S1 + O3S1 -> C10H17N3O9S2 |
| | 3886 | 292.0729 | 12.75 | 4.45 | C10H16N2O6S1 | 0.02 | C10H17N3O6S1 - H1N1 -> C10H16N2O6S1 |
| | 4964 | 306.0522 | 13.52 | 4.8 | C10H14N2O7S1 | -0.09 | C10H17N3O6S1 - H3N1O-1 -> C10H14N2O7S1 |
| B | 115 | 243.0871 | 13.59 | 6.85 | C7H18N1O6P1 | 0.3 | C8H20N1O6P1 - C1H2 -> C7H18N1O6P1 |
| | 117 | 243.0871 | 13.08 | 5.46 | C7H18N1O6P1 | 0.3 | C8H20N1O6P1 - C1H2 -> C7H18N1O6P1 |
| | 2241 | 271.1184 | 12.29 | 4.99 | C9H22N1O6P1 | 0.27 | C8H20N1O6P1 + C1H2 -> C9H22N1O6P1 |
| | 4702 | 325.1653 | 2.09 | 4.08 | C13H28N1O6P1 | 0.38 | C8H20N1O6P1 + C5H8 -> C13H28N1O6P1 |
| C | 1183 | 221.0324 | 7.07 | 5.42 | C10H7N1O5 | 0.1 | C10H7N1O4 + O1 -> C10H7N1O5 |
| | 1991 | 220.0483 | 7.48 | 5.07 | C10H8N2O4 | 0.48 | C10H7N1O4 + H1N1 -> C10H8N2O4 |
| | 1992 | 220.0483 | 7.08 | 5.03 | C10H8N2O4 | 0.48 | C10H7N1O4 + H1N1 -> C10H8N2O4 |
| | 3251 | 204.0534 | 5.6 | 4.32 | C10H8N2O3 | 0.45 | C10H7N1O4 + H1N1O-1 -> C10H8N2O3 |
| | 4144 | 235.0479 | 3.43 | 4.4 | C11H9N1O5 | 0.73 | C10H7N1O4 + C1H2O1 -> C11H9N1O5 |

**Supplementary Figure 2.** Examples of putative metabolites in yeast negative-mode dataset. (A-C) Subnetwork surrounding glutathione (A), glycerophosphocholine (B), and xanthurenic acid (C). (D) Peak properties and annotations for putative metabolites (yellow nodes) in subnetworks (A)-(C).

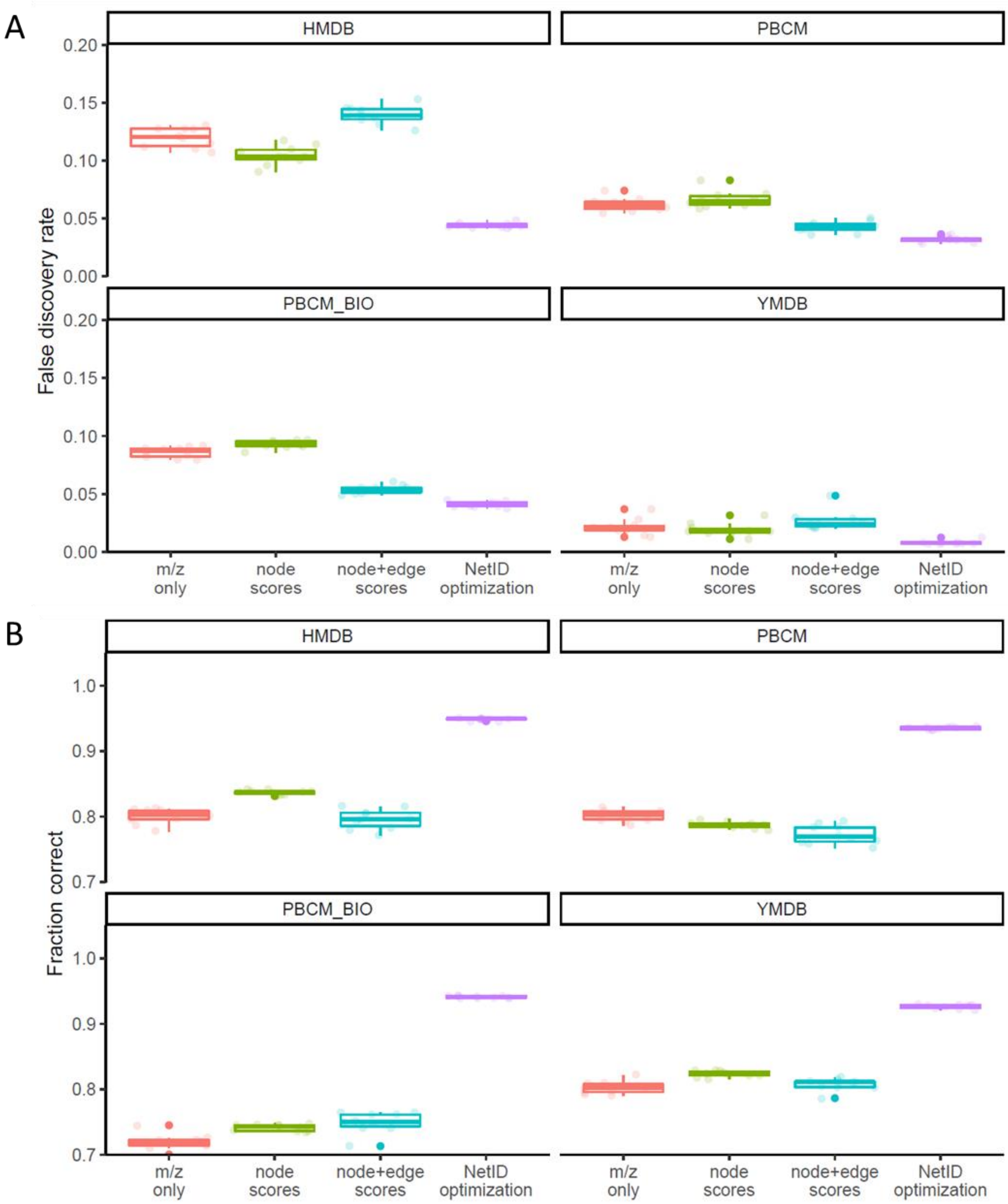


**Supplementary Figure 3.** Evaluation of annotation false discovery rate (FDR) and fraction gold-standard peaks annotated correctly using different reference databases. The four tested reference compound databases are HMDB (human metabolomics database), PBCM (PubChemLite.0.2.0, zenodo.org/record/3611238), PBCM_BIO (a subset of biopathway related entries in PubChemLite.0.2.0) and YMDB (yeast metabolomics database). (A) False discovery rate estimated using target-decoy strategy. Each individual data point (circle) is from a different randomized decoy library (n = 10 randomized libraries were tested for each reference compound database). (B) Fraction of 314 manually curated "ground truth" annotations made correctly (using the 10 randomized libraries for each reference compound database). Boxes show median and interquartile range.

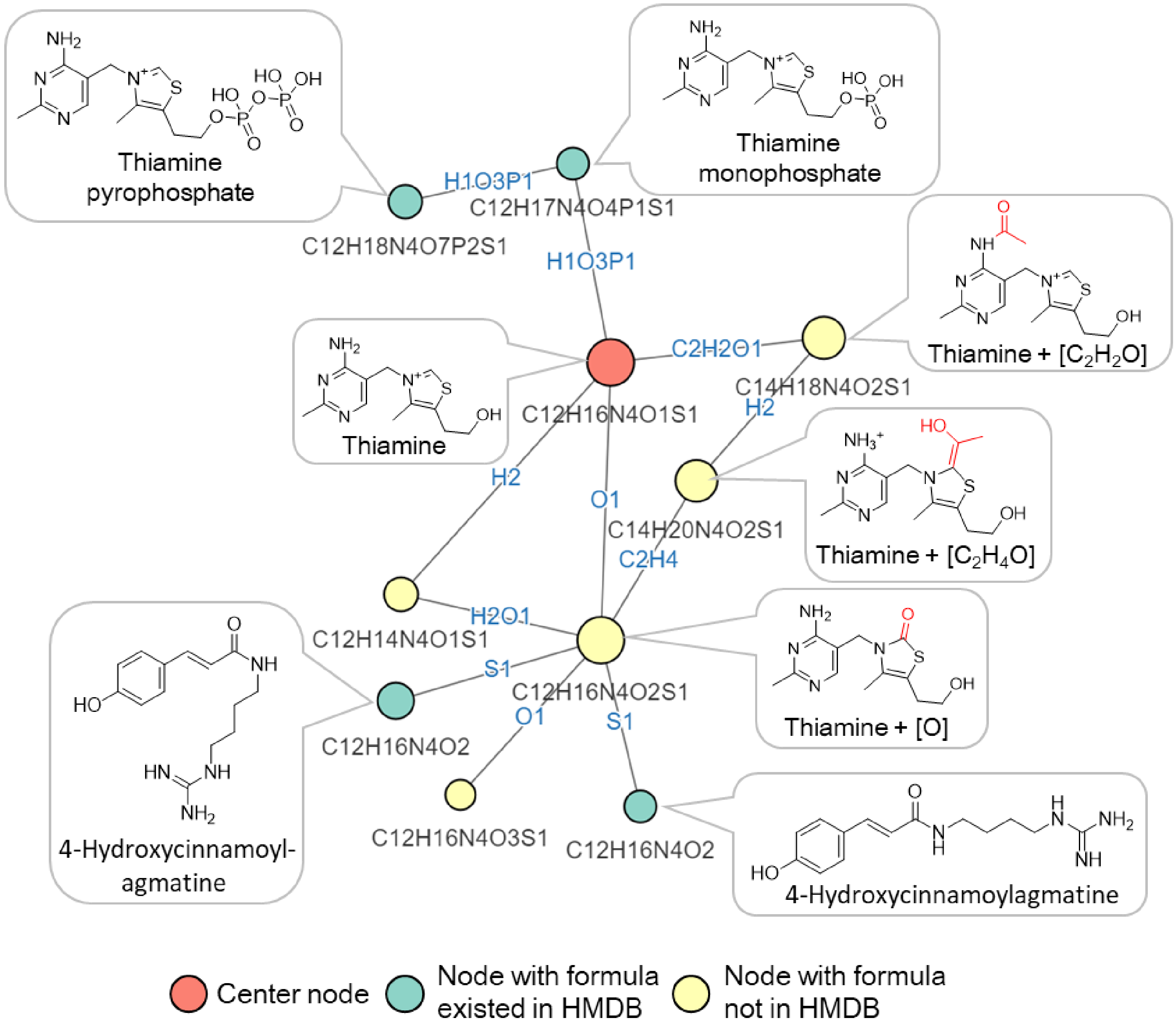


**Supplementary Figure 4.** Subnetwork surrounding thiamine with additional known structures. Nodes, connections, and formulae are direct output of NetID. Boxes with structures were manually added.

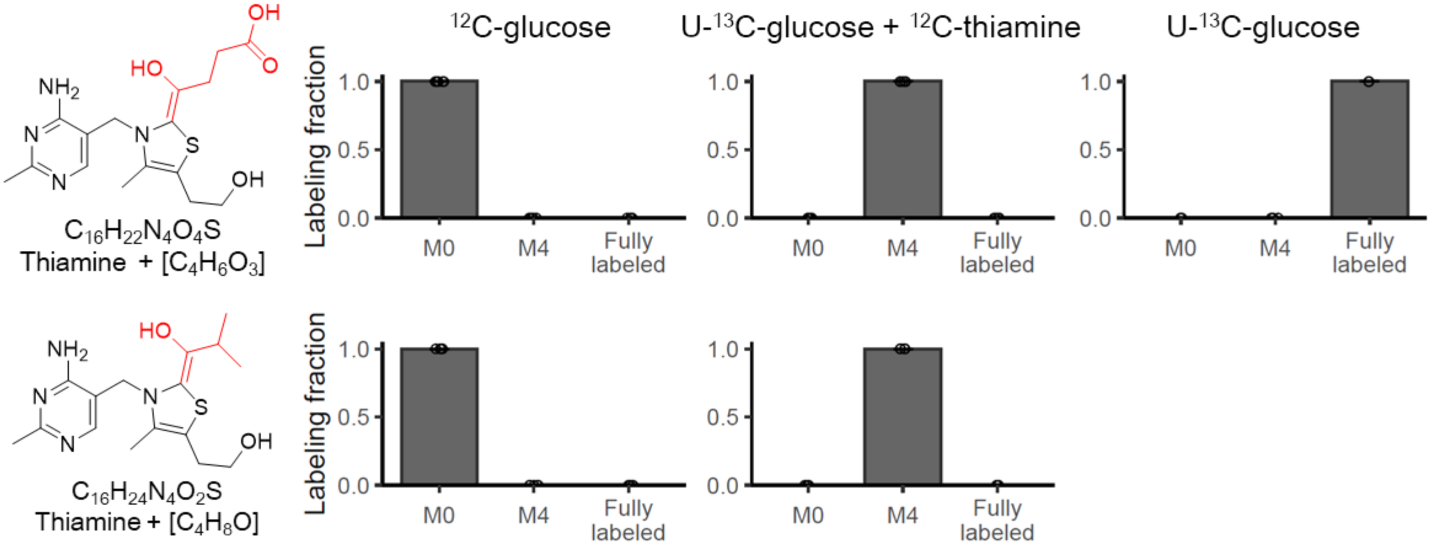


| | M0 | | | M4 | | | Fully labeled | | |
|---|---|---|---|---|---|---|---|---|---|
| | Measured m/z | Expected m/z | Δppm | Measured m/z | Expected m/z | Δppm | Measured m/z | Expected m/z | Δppm |
| Thiamine + [$C_4H_6O_3$] | 367.1439 | 367.1435 | 1.1 | 371.1570 | 371.1569 | 0.3 | 383.1978 | 383.1971 | 1.8 |
| Thiamine + [$C_4H_8O$] | 337.1693 | 337.1693 | 0 | 341.1828 | 341.1827 | 0.3 | | | |

**Supplementary Figure 5.** Evidence for the additional thiamine-derived metabolites. Similar to Figure 3, adding unlabeled thiamine to [U-13C]glucose culture media, yeast uptake the unlabeled thiamine, resulting in unlabeled thiamine, M+4 labeled thiamine+[$C_4H_6O_3$] and thiamine+[$C_4H_8O$] species (n=5). The proposed formulae are also supported by m/z measured by high-resolution mass-spectrometry. Bar represents mean values and error bar indicates s.d..

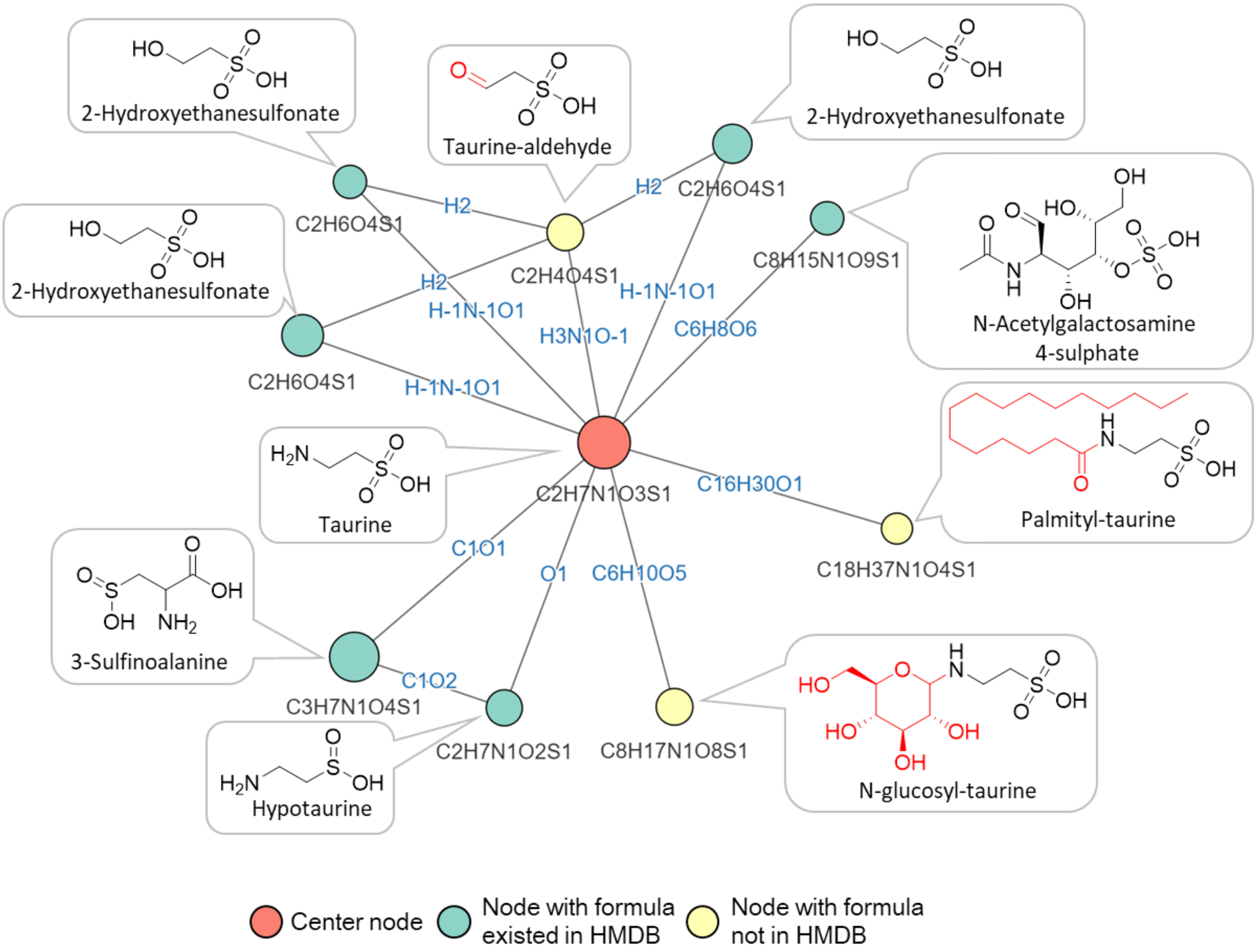


**Supplementary Figure 6.** Subnetwork surrounding taurine with additional known structures. Nodes, connections, and formulae are direct output of NetID. Boxes with structures were manually added.

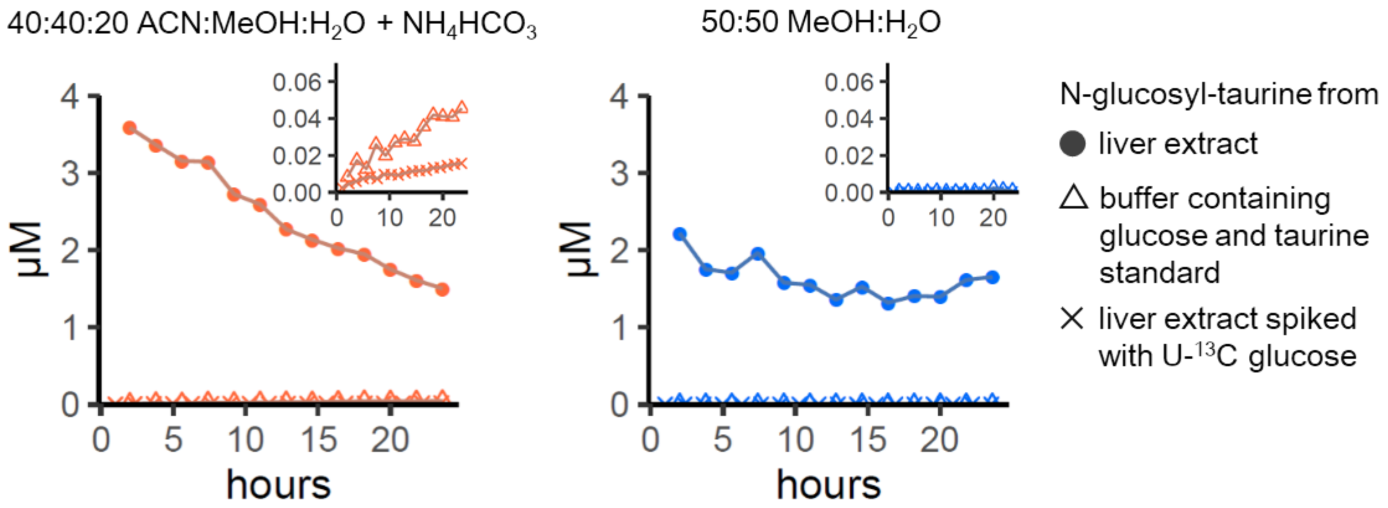


**Supplementary Figure 7.** Glucosyl-taurine is a liver metabolite, not *ex vivo* reaction product. To test for *ex vivo* production of glucosyl-taurine, liver extract (with or without spiked 55 μM [U-$^{13}$C]glucose) or extraction buffer (40:40:20 ACN:MeOH:$H_2O$ + $NH_4HCO_3$ or 50:50 MeOH:$H_2O$) containing pure glucose and taurine were incubated at 5°C for the indicated duration. Metabolites formed by *ex vivo* reactions typically accumulate upon sample incubation, while glucosyl-taurine does not. Moreover, there is minimal assimilation of [U-$^{13}$C]glucose into glucosyl-taurine to make M+6 glucosyl-taurine in liver extract, and, while trace glucosyl-taurine can be formed abiotically in acetonitrile:methanol:water at pH = 7, the observed biological quantity is 100-fold greater.

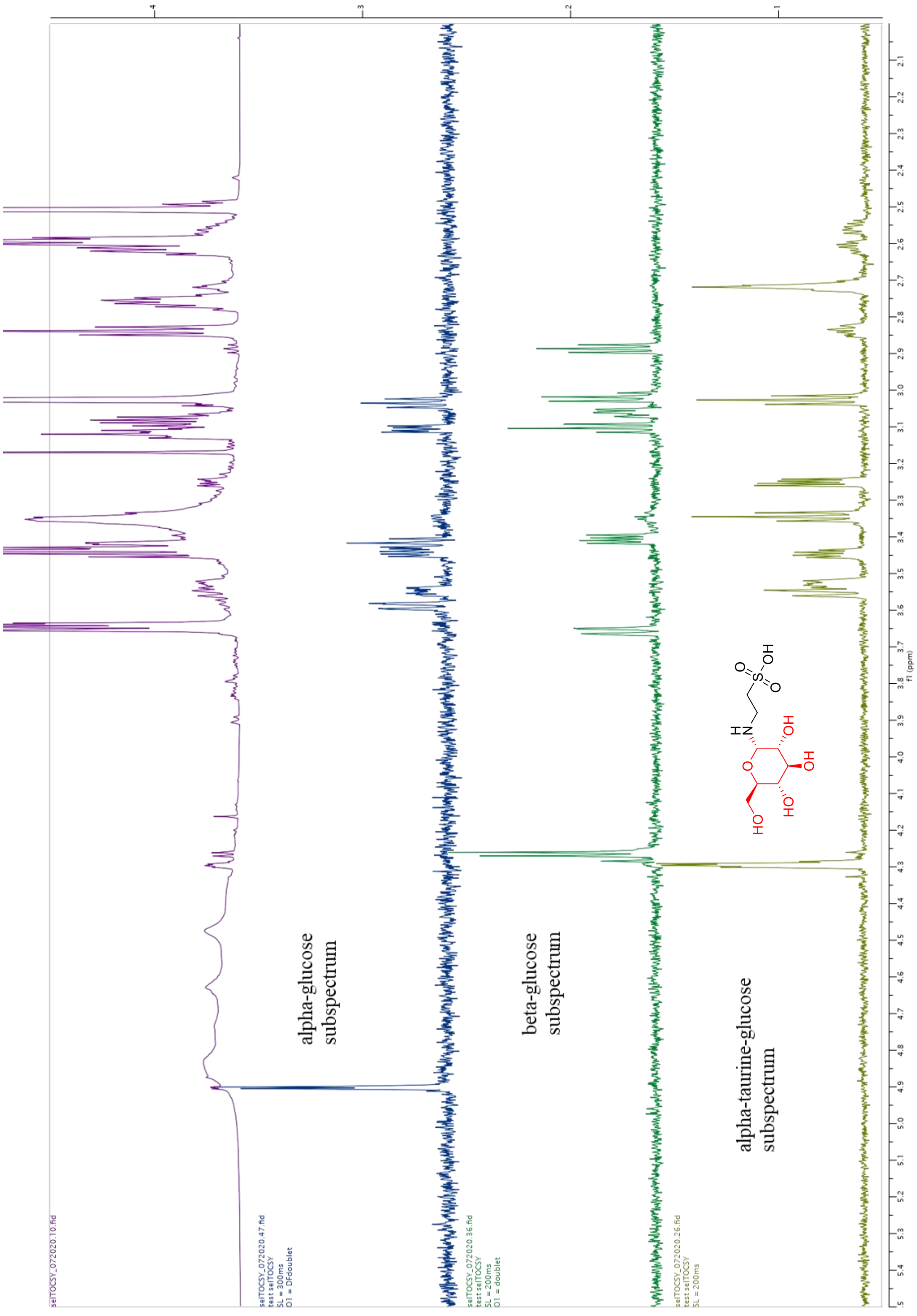


**Supplementary Figure 6.** SelTOCSY NMR confirmation of the structure of the chemically synthesized N-glucosyl-taurine. The final crude material is a mixture of glucose, taurine, and N-glucosyl-taurine at 5.2% (pink line). Comparing N-glucosyl-taurine (yellow) to alpha- (blue) and beta-glucose (green) NMR experiments indicate that C1 of the glucosyl group connects the amine group of taurine in α-position.

**Supplementary table 1.** List of biochemical atom differences

| Type | Formula / Atom difference | Mass difference | RDBE[1] difference | Allowed propagation direction[2] |
|---|---|---|---|---|
| Deamination | O1N-1H-1 | 0.98402 | 0 | 0 |
| Transamination | N1H3O-1 | 1.03163 | -1 | 0 |
| Hydrogenation | H2 | 2.01565 | -1 | 0 |
| Methylation | C1H2 | 14.01565 | 0 | 0 |
| Amination | N1H1 | 15.01090 | 0 | 0 |
| Hydroxylation | O1 | 15.99491 | 0 | 0 |
| Amination | N1H3 | 17.02655 | -1 | 0 |
| Hydration | H2O1 | 18.01056 | -1 | 0 |
| Formylation | C1O1 | 27.99491 | 1 | 0 |
| Beta oxidation | C2H4 | 28.03130 | 0 | 0 |
| Deamination | C1H2O1 | 30.01056 | 0 | 0 |
| Thiolation | S1 | 31.97207 | 0 | 0 |
| Sulfurization | H2S1 | 33.98772 | -1 | 0 |
| Acetylation | C2H2O1 | 42.01056 | 1 | 0 |
| Carboxylation | C1O2 | 43.98983 | 1 | 0 |
| Isoprenylation | C5H8 | 68.06260 | 1 | 1 |
| Sulfurylatoin | S1O3 | 79.95681 | 0 | 0 |
| Phosphorylation | H1P1O3 | 79.96633 | 0 | 0 |
| Hexose | C6H10O5 | 162.05282 | 1 | 1 |
| Uronate | C6H8O6 | 176.03209 | 2 | 1 |
| Palmitoylation | C16H30O1 | 238.22967 | 1 | 1 |
| Sialic acid | C11H17N1O8 | 291.09542 | 3 | 1 |
| AMP | C10H12N5O6P1 | 329.05252 | 7 | 1 |
| CDP | C9H13N3O10P2 | 385.00762 | 5 | 1 |
| ADP-ribosylation | C15H21N5O13P2 | 541.06111 | 8 | 1 |

1. RDBE stands for ring and double bond equivalence.
2. Allowed propagation direction: “1” means only forward propagation is allowed, i.e. adding the indicated atom differences to the parent formula; “-1” means reverse only, i.e. subtracting the indicated atom differences from the parent formula, and “0” means propagation is allowed for both directions.

**Supplementary table 2.** List of abiotic atom differences

| Type | Formula / Atom difference | Mass difference | RDBE difference | Allowed propagation direction |
|---|---|---|---|---|
| Isotope | [10]B-1B1 | 0.99637 | 0 | -1 |
| Isotope | [15]N1N-1 | 0.99703 | 0 | 1 |
| Isotope | [29]Si-1[30]Si1 | 0.99728 | 0 | 1 |
| Isotope | [29]Si1Si-1 | 0.99957 | 0 | 1 |
| Isotope | [53]Cr1Cr-1 | 1.00014 | 0 | 1 |
| Isotope | [13]C1C-1 | 1.00335 | 0 | 1 |
| Isotope | [2]H1H-1 | 1.00628 | 0 | 1 |
| Isotope | [34]S1S-1 | 1.99580 | 0 | 1 |
| Isotope | [30]Si1Si-1 | 1.99684 | 0 | 1 |
| Isotope | [37]Cl1Cl-1 | 1.99705 | 0 | 1 |
| Isotope | [41]K1K-1 | 1.99812 | 0 | 1 |
| Isotope | [18]O1O-1 | 2.00425 | 0 | 1 |
| Isotope | [44]Ca1Ca-1 | 3.99289 | 0 | 1 |
| Isotope | [60]Ni1Ni-1 | 1.99544 | 0 | 1 |
| Isotope | [62]Ni1Ni-1 | 3.99300 | 0 | 1 |
| Adduct | H-1Na1 | 21.98194 | 0 | 1 |
| Adduct | Cl1H1 | 35.97668 | 0 | 1 |
| Adduct | H-1K1 | 37.95588 | 0 | 1 |
| Adduct | H-2Ni1 | 55.91969 | 0 | 1 |
| Adduct | Ca1H-2 | 37.94694 | 0 | 1 |
| Adduct | C1H2O2 | 46.00548 | 0 | 1 |
| Adduct | C1H1Na1O2 | 67.98742 | 0 | 1 |
| Adduct | C1H1K1O2 | 83.96136 | 0 | 1 |
| Adduct | C2H4O2 | 60.02113 | 0 | 1 |
| Adduct | C2H3Na1O2 | 82.00307 | 0 | 1 |
| Adduct | C2H3K1O2 | 97.97701 | 0 | 1 |
| Adduct | C2H2Ni1O2 | 115.94082 | 0 | 1 |
| Adduct | C2Ca1H2O2 | 97.96807 | 0 | 1 |
| Adduct | H2O4S1 | 97.96738 | 0 | 1 |

| Type | Formula / Atom difference | Mass difference | RDBE difference | Allowed propagation direction |
|---|---|---|---|---|
| Adduct | H1Na1O4S1 | 119.94932 | 0 | 1 |
| Adduct | H1K1O4S1 | 135.92326 | 0 | 1 |
| Adduct | H1N1O3 | 62.99564 | 0 | 1 |
| Adduct | N1Na1O3 | 84.97759 | 0 | 1 |
| Adduct | H2C1O3 | 62.00039 | 0 | 1 |
| Adduct | Na1H1C1O3 | 83.98234 | 0 | 1 |
| Adduct | K1H1C1O3 | 99.95628 | 0 | 1 |
| Adduct | H3O4P1 | 97.97690 | 0 | 1 |
| Adduct | H2Na1O4P1 | 119.95884 | 0 | 1 |
| Adduct | H2K1O4P1 | 135.93278 | 0 | 1 |
| Adduct | Cr1O3 | 99.92525 | 0 | 1 |
| Adduct | H4O4Si1 | 95.98789 | 0 | 1 |
| Adduct | H3N1 | 17.02655 | 0 | 1 |
| Adduct | H-2Na2 | 43.96389 | 0 | 1 |
| Adduct | H-2K2 | 75.91176 | 0 | 1 |
| Adduct | C1H1N1 | 27.01090 | 0 | 1 |
| Adduct | C1H4O1 | 32.02621 | 0 | 1 |
| Adduct | H6O8P2 | 195.95379 | 0 | 1 |
| Adduct | B1H-3 | 7.98583 | 2 | 1 |
| Adduct | B1H-1O1 | 25.99640 | 1 | 1 |
| Adduct | H2O3Si1 | 77.97732 | 0 | 1 |
| Adduct | H1Na1O1 | 39.99251 | 0 | 1 |
| Adduct | H1K1O1 | 55.96645 | 0 | 1 |
| Adduct | C2H3N1 | 41.02655 | 0 | 1 |
| Adduct | C3H8O3Si1 | 120.02427 | 0 | 1 |
| Fragment | C1O2 | 43.98983 | 1 | -1 |
| Fragment | C1H2O1 | 30.01056 | 0 | -1 |
| Fragment | H2O1 | 18.01056 | -1 | -1 |
| Fragment | N1H3 | 17.02655 | -1 | -1 |
| Radical | H | 1.00783 | -0.5 | -1 |

**Supplementary table 3.** Examples of nickel related peaks

| peak_id | medMz | medRt | log10_ inten | formula | Δppm | annotation |
|---|---|---|---|---|---|---|
| 4 | 89.0476 | 13.01 | 8.65 | C3H7N1O2 | 0.92 | C3H7N1O2 |
| 1194 | 294.0362 | 13.01 | 5.37 | C8H16N2Ni1O6 | 0.01 | C3H7N1O2 * 2 + C2H2Ni1O2 |
| 1952 | 296.0316 | 13.01 | 4.98 | [60]Ni1C8H16N2O6 | 0.16 | C3H7N1O2 * 2 + C2H2Ni1O2 + [60]Ni1Ni-1* |
| 2723 | 234.0149 | 13 | 4.67 | C6H12N2Ni1O4 | 0.74 | C3H7N1O2 * 2 + H-2Ni1 |
| 2269 | 265.0095 | 13.07 | 4.92 | C7H13N1Ni1O6 | 0.58 | C3H7N1O2 + C2H4O2 + C2H2Ni1O2 |
| 3869 | 267.005 | 13.04 | 4.36 | [60]Ni1C7H13N1O6 | 0.36 | C3H7N1O2 + C2H4O2 + C2H2Ni1O2 + [60]Ni1Ni-1 |
| 855 | 270.0961 | 13.59 | 5.62 | C10H14N4O5 | 1.27 | C10H14N4O5 |
| 2005 | 358.0423 | 13.5 | 4.97 | C11H16N4Ni1O6 | 0.14 | C10H14N4O5 + C1H4O1 + H-2Ni1 |
| 2961 | 360.0377 | 13.51 | 4.39 | [60]Ni1C11H16N4O6 | 0.26 | C10H14N4O5 + C1H4O1 + H-2Ni1 + [60]Ni1Ni-1 |
| 125 | 222.0673 | 14.18 | 6.82 | C7H14N2O4S1 | 0.68 | C7H14N2O4S1 |
| 2406 | 338.0083 | 14.29 | 4.95 | C9H16N2Ni1O6S1 | -0.08 | C7H14N2O4S1 + C2H2Ni1O2 |
| 31 | 117.079 | 11.21 | 7.61 | C5H11N1O2 | 0.02 | C5H11N1O2 |
| 3450 | 293.041 | 11.24 | 4.54 | C9H17N1Ni1O6 | -0.16 | C5H11N1O2 + C2H4O2 + C2H2Ni1O2 |
| 4664 | 146.069 | 13.39 | 8.28 | C5H10N2O3 | 1.13 | C5H10N2O3 |
| 3287 | 262.0098 | 13.4 | 4.53 | C7H12N2Ni1O5 | 0.72 | C5H10N2O3 + C2H2Ni1O2 |
| 3534 | 322.0311 | 13.37 | 4.38 | C9H16N2Ni1O7 | 0.05 | C5H10N2O3 + C2H2Ni1O2 + C2H4O2 |
| 92 | 612.1521 | 14.19 | 6.98 | C20H32N6O12S2 | -0.19 | C20H32N6O12S2 |
| 2440 | 668.0718 | 14.2 | 4.87 | C20H30N6Ni1O12S2 | -0.18 | C20H32N6O12S2 + H-2Ni1 |
| 3528 | 670.0671 | 14.21 | 4.54 | [60]Ni1C20H30N6O12S2 | 0.03 | C20H32N6O12S2 + H-2Ni1 + [60]Ni1Ni-1 |
| 22 | 132.0898 | 16.24 | 7.34 | C5H12N2O2 | 0.77 | C5H12N2O2 |
| 4942 | 308.052 | 16.26 | 4.1 | C9H18N2Ni1O6 | -0.48 | C5H12N2O2 + C2H4O2 + C2H2Ni1O2 |

* "[60]Ni1Ni-1" means adding an isotope $^{60}$Ni and subtracting a regular Ni, representing the atom difference of nickel isotope. This representation aligns atom difference to mass difference.

**Supplementary table 4.** Search results of reported novel metabolites in compound databases.

| | HMDB | | PubChem | | METLIN | | ChemSpider | |
|---|---|---|---|---|---|---|---|---|
| | formula | structure | formula | structure | formula | structure | formula | structure |
| Thiamine + [$C_2H_2O$] ($C_{14}H_{18}N_4O_2S$) | X | X | √ | X | X | X | √ | X |
| Thiamine + [$C_2H_4O$] ($C_{14}H_{20}N_4O_2S$) | X | X | √ | X[1] | X | X | √ | X[1] |
| Thiamine + [$C_4H_6O_3$] ($C_{16}H_{22}N_4O_4S$) | X | X | √ | X[1] | X | X | √ | X[1] |
| Thiamine + [$C_4H_8O$] ($C_{16}H_{24}N_4O_2S$) | X | X | √ | X[1] | X | X | √ | X[1] |
| Glucosyl-taurine ($C_8H_{17}NO_8S$) | X | X | √ | √[2] | X | X | √ | X |

**Note:**
1. Pyrophosphate form of the metabolite exists.
2. Reported only as a synthetic chemical, not a metabolite or biological chemical.

**Supplementary Table 5.** Memory and run-time used in NetID.

| | Yeast neg | Yeast pos | Liver neg | Liver pos |
|---|---|---|---|---|
| Total non-background peaks | 5588 | 9833 | 8191 | 12128 |
| Maximum memory used (GB) | 4.7 | 13.3 | 6.8 | 12.1 |
| Optimization time (min) | 2.5 | 6.7 | 1.1 | 2.4 |
| Total time (min) | 24.3 | 101.3 | 31.8 | 94.4 |

**Note:** The maximum memory and run-time reported here is under default parameter setting in NetID.

### Supplementary Note 1 - NetID User Guide

***Li Chen, Ziyang Chen***

**6/1/2021**

A primary goal of liquid chromatography-high resolution mass spectrometry (LC-MS)-based metabolomics is to identify all metabolites, but most LC-MS peaks remain unidentified. Here, we present a global network optimization approach, NetID, to annotate untargeted LC-MS metabolomics data. We consider all experimentally observed ion peaks together, and assign annotations to all of them simultaneously so as to maximize a score that considers properties of peaks (known masses, retention times, MS/MS fragmentation patterns). Global optimization results in accurate peak assignment and trackable peak-peak relationships in the output network. Applying this approach to yeast and mouse data, we identify 5 novel metabolites, including thiamine and taurine derivatives. Isotope tracer studies indicate active flux through these metabolites. Thus, NetID applies existing metabolomic knowledge and global optimization to annotate untargeted metabolomics data, revealing novel metabolites.

NetID requires (1) a peak table containing m/z, RT and intensity from high-resolution mass spectrometry data. (2) a reference compound database, which we provide HMDB, YMDB, a lite version of PubChem (PubChemLite.0.2.0) and a subset of 47,101 biopathway related entries (PubChemLite_Bio) for user to choose. (3) a transformation table, which we assembled a list of 25 biochemical atom differences and 59 abiotic atom differences. NetID optionally use (4) a list of excel files containing MS2 fragmentation information (m/z and intensity) for peaks in the above peak table. (5) a list of known metabolites' retention time, which we provide our in-house retention time list for demonstration. Users can customize the compound database, the transformation table and the retention time list following the user guide. Currently the algorithm is developed using Thermo Orbitrap instruments results. We anticipate the algorithm will work for other high mass accuracy data, such as TOF data, but parameters may need to be optimized for the best performance.

Citation: https://www.biorxiv.org/content/10.1101/2021.01.06.425569v1
Git-hub: https://github.com/LiChenPU/NetID

### 1 Environment Setup

This section provides step-by-step instructions to set up the environment to run NetID algorithm in a local computer. A Windows system is recommended. Typical install time on a "normal" desktop computer is within a few hours.

#### 1.1 Software installation

- Install R, Rstudio, Rtools40, ILOG CPLEX Optimization Studio (CPLEX), preferably at default location.

**R**(4.0 or later): https://www.r-project.org/
**RStudio**: https://rstudio.com/products/rstudio/download
**Rtools40**: https://cran.r-project.org/bin/windows/Rtools/ow
**CPLEX**(12.8 or later): https://www.ibm.com/academic/technology/data-science

- You need to add R and Rtools40 to Environmental Variables PATH, with instruction provided at the end.

#### 1.2 Code download

##### 1.2.1 Via Git (recommended)

1. Install **git** via https://support.rstudio.com/hc/en-us/articles/200532077?version=1.3.1093&mode=desktop
2. In Rstudio, go to `File` → `New project` → `Version control` → `Git`, enter https://github.com/LiChenPU/NetID.git for `URL`, select a subdirectory, and create project.
3. You should be able to see all files in place under your selected subdirectory. Use pull option to check for latest updates.

##### 1.2.2 Via Github

1. Go to website https://github.com/LiChenPU/NetID, hit the green `code` button, select download zip, and unzip files.

#### 1.3 Package dependency installation

Most of the dependent packages can be installed by running the R script `NetID_packages.R` in the `get started` folder. See **Troubleshooting** section for possible errors.

The package, **cplexAPI**, connecting R to CPLEX, requires additional installation steps.

1. Go to website: https://cran.r-project.org/web/packages/cplexAPI/index.html, look for `Package source`, and download `cplexAPI_1.4.0.tar.gz`. In the same page, look for `Materials`, a package installation guide can be found in the link `INSTALL`.
2. Unzip the folder `cplexAPI` to the **desktop**, open subfolder `src`, follow the installation guide to modify the file `Makevars.win`.
   *Note*: Replace `\` in the `Makevars.win` file into `/` in order for R to recognize the path.
   - For example, the `-I"${CPLEX_STUDIO_DIR}\cplex\include"` should be replaced with the path CPLEX_studio is installed, such as:
     `-I"C:/Program Files/IBM/ILOG/CPLEX_Studio1210/cplex/include"`
   - The `-L"${CPLEX_STUDIO_LIB}"` should be replaced with the path CPLEX_studio is installed, such as:
     `-L"C:/Program Files/IBM/ILOG/CPLEX_Studio1210/cplex/bin/x64_win64"`
3. In command line, run line below to build package, change `${Username}` to actual name.
   `R CMD build --no-build-vignettes --no-manual --md5 "C:\Users\${Username}\Desktop\cplexAPI"`
   a new package `cplexAPI_1.4.0.tar.gz` will be built under the default path (for example, `C:\Users\${Username}`)

4. In command line, run line below to install package.
   `R CMD INSTALL --build --no-multiarch .\cplexAPI_1.4.0.tar.gz`
   If you see `DONE (cplexAPI)`, then the package installation is successful.

   - *Note*: if error occurs relating to `__declspec(dllimport deprecated)`, you need to go to `C:\Program Files\IBM\ILOG\CPLEX_Studio1210\cplex\include\ilcplex` (or other installation path), open the `file cpxconst.h`, go to the line indicated in the error message or search for `__declspec(dllimport deprecated)`, add `_` to `__declspec(dllimport deprecated)`, making it to `__declspec(dllimport_deprecated)`. Save file and repeat *step 4*.

5. To take a short venture using CPLEX in R, refer to **Package cplexAPI – Quick Start** in https://cran.r-project.org/web/packages/cplexAPI/index.html.

### 2 Using NetID

This section will use yeast negative-mode dataset and mouse liver negative-mode dataset as examples to walk through the NetID workflow.

- *Note 1: If other El-MAVEN version was used, check the "raw_data.csv" for the column number where the first sample is located, and specify that in the NetID_run_script.R file. For example, In El-MAVEN (version 7.0), first_sample_col_num is set at 15 as default. If El-MAVEN (version 12.0) is used, first_sample_col_num should be set at 16.*
- *Note 2: for more advanced uses, scoring and other parameters can be edited in NetID_function.R and NetID_run_script.R. Read the manuscript method section for detailed explanation on parameters.*

#### 2.1 Yeast negative-mode dataset

In the `Sc_neg` folder, file `raw_data.csv` is the output from **Elmaven** recording MS information, and is the input file for **NetID**. MS2 is not collected for this dataset.

##### 2.1.1 Running the code

1. Open `code folder` → `NetID_run_script.R`
2. In the `# Setting path #### section`, set work_dir as `"../Sc_neg/"`.

```
# Setting path ####
{
  setwd(dirname(rstudioapi::getSourceEditorContext()$path))
  source("NetID_function.R")

  work_dir = "../Sc_neg/"
  setwd(work_dir)
  printtime = Sys.time()

}
```

3. In the `# Read data and files #### section`, set filename as `"raw_data.csv"`, set MS2_folder as `""`.
   set `ion_mode` as `-1` if negative ionization data is loaded, and `1` if positive ionization data loaded.

```
# Read data and files ####
{
  Mset = list()
  # Read in files
  Mset = read_files(filename = "raw_data.csv",
                    LC_method = "Hilic_25min_QE",
                    ion_mode = -1 # 1 for pos mode and -1 for neg mode
                    )
  Mset = read_MS2data(Mset,
                      MS2_folder = "") # MS2
}
```

4. Keep all other parameters as default, and run all lines.

##### 2.1.2 Expected outputs

1. In the console, error message should not occur. If optimization step is successful, you will see messages in the following format.

```
"Optimization ended successfull - integer optimal, tolerance - OBJ_value = 2963.71
(bestobjective - bestinteger) / (1e-10 + |bestinteger|) = 0.000048268"
95.74 sec elapsed
```

2. Three files will be generated in the `Sc_neg` folder. Expected run time on a "normal" desktop computer should be within an hour.
   - `NetID_output.csv` contains the annotation information for each peak.
   - `NetID_output.RData` contains node, edge and network information. The file will be used for network visualization in Shiny R app.
   - `.RData` records the environmental information after running codes. The file is mainly used for development and debugging.

#### 2.2 Your own dataset

##### 2.2.1 MS1 dataset preparation

1. File conversion. Use software **ProteoWizard40** (version 3.0.11392) to convert LC-MS raw data files (.raw) into mzXML format. A command line script specifies the conversion parameter. Assuming the raw data are in `D:/MS data/test`. Type in the scripts below.

```
D:
cd D:/MS data/test
"C:/Program Files/ProteoWizard/ProteoWizard 3.0.11392/msconvert.exe"
*.raw --filter "peakPicking true 1-" --simAsSpectra --srmAsSpectra --mzXML
```

If **ProteoWizard** is installed in location other than `C:/Program Files/ProteoWizard/ProteoWizard 3.0.11392/msconvert.exe`, specify your path to where you can find the `msconvert.exe` file. Expected outputs will be `.mzXML` files from `.raw` data.

2. **El-MAVEN (version 7.0)** is used to generate a peak table containing m/z, retention time, intensity for peaks. Detailed guides for peak picking can be found in https://elucidatainc.github.io/ElMaven/faq/.
   After peak picking and a peak table tab has shown up, click `export to CSV`. Choose `export all groups`. In the pop-up saved window, choose format `Groups Summary Matrix Format Comma Delimited`. Save to the desired path.
3. Under the `NetID` folder, create a new folder `NetID_test`, copy the csv file from *step 2* into the folder, and change the filename into `raw_data.csv`.

##### 2.2.2 MS2 dataset preparation

**NetID** currently utilizes targeted MS2 data for better MS2 quality, and will incorporate data-dependent MS2 data in the future.

1. Prepare MS2 inclusion list
   For targeted MS2 analysis, from the peak list generated in *step 1*, select the peaks (m/z, RT) that you want to perform MS2, and arrange them into multiple csv files that will serve as the inclusion lists to set up the PRM method on **Thermo QExactive** instrument. Instruction can be found in https://proteomicsresource.washington.edu/docs/protocols05/PRM_QExactive.pdf.
   *Note*: Arrange the parent ions so as to avoid to perform many PRMs at same time. An example is shown below with the start and End time set as RT-1.5 and RT+1.5 (min) to have good chromatogram coverage.

```
library(readr)
read_csv("example.csv")
##
## -- Column specification --------------------------------------------------------
## cols(
##   Mass = col_double(),
##   Formula = col_logical(),
##   Formula_type = col_logical(),
##   Species = col_logical(),
##   CS = col_logical(),
##   Polarity = col_character(),
##   Start = col_double(),
##   End = col_double(),
##   CE = col_double(),
##   CE_type = col_character(),
##   MSXID = col_logical(),
##   Comment = col_character()
## )
## # A tibble: 16 x 12
##      Mass Formula Formula_type Species CS    Polarity Start   End    CE CE_type
##     <dbl> <lgl>   <lgl>        <lgl>   <lgl> <chr>    <dbl> <dbl> <dbl> <chr>
##   1 499.  NA      NA           NA      NA    Negative 0.456  3.46    30 NCE
##   2 722.  NA      NA           NA      NA    Negative 0.733  3.73    30 NCE
##   3 403.  NA      NA           NA      NA    Negative 1.06   4.06    30 NCE
```

```
##  4 211.  NA      NA          NA      NA    Negative  1.20   4.20    30 NCE
##  5 328.  NA      NA          NA      NA    Negative  1.40   4.40    30 NCE
##  6 149.  NA      NA          NA      NA    Negative  1.59   4.59    30 NCE
##  7 151.  NA      NA          NA      NA    Negative  2.69   5.69    30 NCE
##  8 335.  NA      NA          NA      NA    Negative  2.70   5.70    30 NCE
##  9 143.  NA      NA          NA      NA    Negative  4.07   7.07    30 NCE
## 10  89.0 NA      NA          NA      NA    Negative  5.67   8.67    30 NCE
## 11 283.  NA      NA          NA      NA    Negative  6.92   9.92    30 NCE
## 12 202.  NA      NA          NA      NA    Negative  8.79  11.8     30 NCE
## 13 160.  NA      NA          NA      NA    Negative 10.3   13.3     30 NCE
## 14 216.  NA      NA          NA      NA    Negative 11.4   14.4     30 NCE
## 15 125.  NA      NA          NA      NA    Negative 12.0   15.0     30 NCE
## 16 230.  NA      NA          NA      NA    Negative 12.9   15.9     30 NCE
## # ... with 2 more variables: MSXID <lgl>, Comment <chr>
```

2. Instrument setup
   Set up the **QExactive** instrument so that it contains both "Full MS" and "PRM" scan events. For PRM setup, use the above file as inclusion list to perform targeted MS2 analysis. We typically use the following setting for MS2 analysis: resolution 17500, AGC target 1e6, Maximum IT 500 ms, isolation window 1.5 m/z. For a total of 1500 parent ions and 15 parent ions for each method, it requires a total of 100 runs, or ~42 hours using a 25-min LC method.

3. MS2 file conversion.
   **RawConverter** (version 1.2.0.1, http://fields.scripps.edu/rawconv/) is used to convert the `.raw` file into `.mzXML` file that contains MS2 information. Keep the default parameters except setting `Environment Type` as `Data Independent`, and `Output Formats` as `mzXML`.

4. MS2 reading and cleaning.
   A matlab code is used for MS2 reading and cleaning, which can be found in **CodeOcean** as a published capsule (https://codeocean.com/capsule/1048398/tree/v1). The `csv` files from *1* paired with the MS2 data files in `mzXML` format from *3* are the required input data. Refer to capsule description and `readme.md` file for more details of how the code works. In Brief,

   - Prepare filename. Filenames for both csv and mzXML files should be named as `prefixNNN`, where prefix is the given file name and NNN is the 3 digits number in continuous order (e.g. `M001.csv`, `M002.csv`,... and `M001.mzXML`, `M002.mzXML`,... in the `/data` folder).

   - Duplicate the capsule to your own account so you can edit and use the capsule. Upload your own files and remove the previous files in `/data` folder.

   - Specify the prefix and the range of numbers at the beginning section of the main code `Main_example.m`.

   - Set the main code as file to run in Code Ocean using the dropdown menu next to main code.

   - Click `reproducible run` to perform the batch processing.

- The resulting output files in `.xlsx` format with the same filenames will appear in the timeline. Each `xlsx` file contains multiple tabs of cleaned MS2 spectra. The names of the tabs correspond to the row numbers of the `csv` file specifying the individual parent peak information.

5. Save files to folders.
   Back to the `NetID_test` folder, create a new folder `MS2`, download all `xlsx` files from *4* into the folder.

##### 2.2.3 Running the code

1. Open `code` folder → `NetID_run_script.R`.
2. In the `# Setting path #### section`, set `work_dir` as `"../NetID_test/"`.
3. In the `# Read data and files ####` section,
   set `filename` as `raw_data.csv`, set `MS2_folder` as `MS2`.
   set `LC_method` to specify column to read for the retention time of known standards. (In folder `NetID` → `dependent` → `known_library.csv`, update the retention time info as needed.)
   set `ion_mode` as `-1` if negative ionization data is loaded, and `1` if positive ionization data loaded.
4. Keep all other parameters as default, and run all lines.

##### 2.2.4 Expected outputs

Similar to the `demo` file, the console will print out message indicating optimization step is successful, and three files `NetID_output.csv`, `NetID_output.RData` and `.RData` will be generated in the `NetID_test` folder

#### 2.3 Other Settings

##### 2.3.1 Compound librarys

###### 2.3.1.1 Other provided librarys

**NetID** provides 4 librarys for the user to choose: **HMDB**, **YMDB**, **PubChem**, **PubChem Bio-pathway only**.

To select the desired database, change `HMDB_library_file = "../dependent/hmdb_library.csv"` to `../dependent/ymdb_library.csv`, `../dependent/pbcm_library.csv` or `../dependent/pbcm_library_bio.csv`.

```
Mset = read_files(filename = "raw_data.csv",
                  LC_method = "Hilic_25min_QE",
                  ion_mode = -1, # 1 for pos mode and -1 for neg mode
                  HMDB_library_file = "../dependent/hmdb_library.csv"
                  )
```

###### 2.3.1.2 Design your own library

1. A workable library requires following columns.

```
read_csv("../../dependent/hmdb_library.csv")
##
```

```
## -- Column specification --------------------------------------------------------
## cols(
##   accession = col_character(),
##   iupac_name = col_character(),
##   name = col_character(),
##   SMILES = col_character(),
##   status = col_character(),
##   formula = col_character(),
##   mass = col_double(),
##   rdbe = col_double(),
##   category = col_character()
## )
## # A tibble: 114,014 x 9
##    accession  iupac_name     name  SMILES    status formula  mass  rdbe category
##    <chr>      <chr>          <chr> <chr>     <chr>  <chr>   <dbl> <dbl> <chr>
##  1 HMDB00000~ (2S)-2-amino-~ 1-Me~ CN1C=NC(~ quant~ C7H11N~ 169.      4 Metabol~
##  2 HMDB00000~ propane-1,3-d~ 1,3-~ NCCCN     quant~ C3H10N2  74.1     0 Metabol~
##  3 HMDB00000~ 2-oxobutanoic~ 2-Ke~ CCC(=O)C~ quant~ C4H6O3  102.      2 Metabol~
##  4 HMDB00000~ 2-hydroxybuta~ 2-Hy~ CCC(O)C(~ quant~ C4H8O3  104.      1 Metabol~
##  5 HMDB00000~ (1S,10R,11S,1~ 2-Me~ [H][C@@]~ quant~ C19H24~ 300.      8 Metabol~
##  6 HMDB00000~ (3R)-3-hydrox~ (R)-~ C[C@@H](~ quant~ C4H8O3  104.      1 Metabol~
##  7 HMDB00000~ 1-[(2R,4S,5R)~ Deox~ OC[C@H]1~ quant~ C9H12N~ 228.      5 Metabol~
##  8 HMDB00000~ 4-amino-1-[(2~ Deox~ NC1=NC(=~ quant~ C9H13N~ 227.      5 Metabol~
##  9 HMDB00000~ (1S,2R,10R,11~ Cort~ [H][C@@]~ quant~ C21H30~ 346.      7 Metabol~
## 10 HMDB00000~ (1S,2R,10S,11~ Deox~ [H][C@@]~ quant~ C21H30~ 330.      7 Metabol~
## # ... with 114,004 more rows
```

2. To build your own library, make a csv file in the same format as the one shown above, and set `HMDB_library_file = "../dependent/hmdb_library.csv"` to your desired directory in `NetID_run_script.R`.

##### 2.3.2 Modifying `emperical_rules.csv`

`emperical_rules.csv` can also be created or modifed to support specific biotransformation. A workable emperical_rules requires following columns. * `name` and `note` is not necessary. * `category` includes: `Biotransform`, `Natural_abundance`, `Adduct`, `Fragment` and `Radical` * `rbde` is calculated using the `formula_rbde` function of the package `lc8` * `direction` states the possible direction of transformation: `1` means from larger mass to smaller mass; `0` means the opposite; `-1` means both direction are possible.

```
read_csv("../../dependent/empirical_rules.csv")
##
## -- Column specification --------------------------------------------------------
## cols(
##   category = col_character(),
##   name = col_character(),
##   formula = col_character(),
##   mass = col_double(),
##   direction = col_double(),
##   rdbe = col_double(),
##   note = col_character()
```

```
## )
## # A tibble: 84 x 7
##    category     name  formula    mass direction  rdbe note
##    <chr>        <chr> <chr>     <dbl>     <dbl> <dbl> <chr>
##  1 Biotransform O-HN  O1N-1H-1  0.984         0     0 Deamination
##  2 Biotransform NH3-O N1H3O-1   1.03          0    -1 Transamination
##  3 Biotransform H2    H2        2.02          0    -1 Hydrogenation
##  4 Biotransform CH2   C1H2     14.0           0     0 Methylation
##  5 Biotransform NH    N1H1     15.0           0     0 Amination
##  6 Biotransform O     O1       16.0           0     0 Hydroxylation
##  7 Biotransform N1H3  N1H3     17.0           0    -1 Amination (+NH3)
##  8 Biotransform H2O   H2O1     18.0           0    -1 Hydration
##  9 Biotransform CO    C1O1     28.0           0     1 Formylation (+CO)
## 10 Biotransform C2H4  C2H4     28.0           0     0 Beta oxidation
## # ... with 74 more rows
```

#### 2.3.3 Retention time list

##### 2.3.3.1 Customize your own RT table

1. In the dependent folder, open the `known_library_customized.csv` file

```
read_csv("../../dependent/known_library_customized.csv")[1:5,]
##
## -- Column specification --------------------------------------------------------
## cols(
##   name = col_character(),
##   HMDB = col_character(),
##   formula = col_character(),
##   SMILES = col_character(),
##   Hilic_25min_QE = col_double(),
##   No_RT = col_logical()
## )
## # A tibble: 5 x 6
##   name                      HMDB     formula    SMILES       Hilic_25min_QE No_RT
##   <chr>                     <chr>    <chr>      <chr>                 <dbl> <lgl>
## 1 1-Methyl imidozolacetic~ HMDB000~ C6H8N2O2   CN1C=C(N=C1~           9.05 NA
## 2 5-L-Hydroxytryptophan    <NA>     C11H12N2O3 <NA>                  10.2  NA
## 3 ADP                      <NA>     C10H15N5O~ <NA>                  13.9  NA
## 4 CDP                      <NA>     C9H15N3O1~ <NA>                  NA    NA
## 5 CDP-choline              <NA>     C14H26N4O~ <NA>                  NA    NA
```

2. Column `Name`, `formula` are required. Column `HMDB`, `SMILES` are optional. For each RT list (e.g. Hilic_25min_QE), record the retention time under the column. Multiple RT lists can be stored by adding additional columns. Empty retention time is allowed for a entry.

**2.3.3.2 Skip RT table** Setting the LC_method = "No_RT". Then RT information will not be considered in the algorithm.

```
Mset = read_files(filename = "raw_data.csv",
                  LC_method = "No_RT",
                  ion_mode = -1, # 1 for pos mode and -1 for neg mode
                  HMDB_library_file = "../dependent/hmdb_library.csv"
                  )
```

##### 2.3.4 Score Setting

See Supplementary Note 2 of NetID paper for explanation

### 3 NetID Visualization

This section provides instruction to visualize and explore **NetID** output results in either **Cytoscape** software or interactive **Shiny R app**. After running **NetID** algorithm, it will export one `.R` and two `.csv`files (`cyto_node.csv` and `cyto_edges.csv`), storing the nodes and edges of the output network.

#### 3.1 Cytoscape

0. What is **Cytoscape** For more info regarding what is **Cytoscape**, check https://cytoscape.org/what_is_cytoscape.html.

1. install **Cytoscape** Download **Cytoscape** (https://cytoscape.org/download.html) and follow installation instruction to install onto your computer.

2. Load the example **NetID** output into **Cytoscape**

   - Run **Cytoscape**, click `import network from file system`, and load `cyto_edges.csv`, set `edge_id` column as the `key`, set `node1`as `source node`, set `node2` column as `target node`, and the rest columns as `edge attribute`.

   - Click `import table from file`, load `cyto_node.csv`, set `node_id` column as the `key`, and the rest columns as `node attribute`.
   - Select `subnetwork`, set `styles`, and explore the network with various functionalities inside **Cytoscape**.

3. Explore in Cytoscape
   http://manual.cytoscape.org/en/stable/index.html provides all you need to know about exploring in **Cytoscape**. (This writer knew little about this cool software, so all he could give was this link and *may the Force be with you*.)

4. Export
   The network as well as the curated subnetworks can be exported for future analysis or sharing with others. An example network file `example.cys` is included along with the two `.csv` files, which is created using **Cytoscape** version 3.8.2

#### 3.2 Shiny App

This part provides instruction to visualize and explore **NetID** output results in the interactive **Shiny R app**. A 21-inch or larger screen is recommended for best visualization.

##### 3.2.1 Runing Shiny App

1. Open `code` folder → `R_shiny_App.R`.
2. In the `# Read in files #### section`, set datapath as `../Sc_neg/`
3. Keep all other parameters as default, and run all lines.
4. A Shiny app will pop up.

##### 3.2.2 Searching peaks of interest

1. On the left panel, you can enter a m/z or a formula to search your peak of interest. For example, 180.0631 or C6H12O6 will automatically update the data table on the right. Enter 0 to restore full list for the data table.
2. Change ionization and ppm window to adjust calculated m/z. W
3. On the right, you can explore the peak list in an interactive data table, including global text search on top right, specifying ranges for numeric column or searching text within character columns, ranking each column etc.

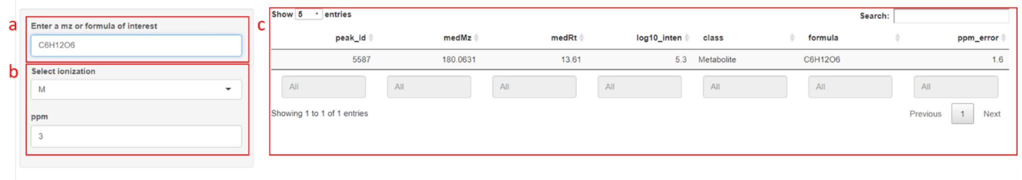


##### 3.2.3 Network Visualization

1. Peak ID, formula and class determines the center node for the network graph. Peak ID will be automatically updated by the first line in the data table if a m/z or formula is given. Alternatively, you can manually enter Peak ID.
2. The degree parameter controls how far the network expands from the center node. Degree 1 means only nodes directly connected to the center node will be shown and degree 2 means nodes connected to degree 1 will be shown, etc.
3. Biochemical graph shows biochemical connections. Abiotic graph shows abiotic connections. Node labels and Edge labels determines if the graph show node or edge labels. Optimized only determines whether to show only the optimal annotations or all possible annotations in the network.
4. When setting parameters, hit plot to see the network graph.

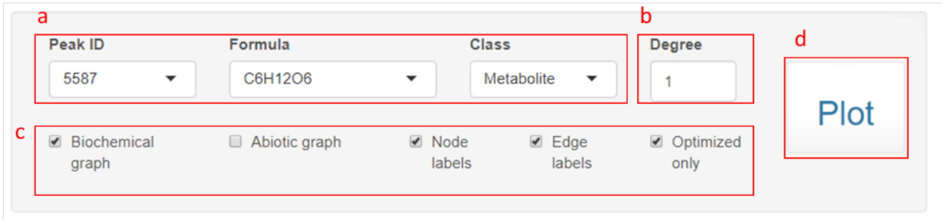


5. A sample network graph is shown below (a different center node may give less complicated graph). You may edit the nodes or edges (top left), move figures with the arrow buttons (bottom left), and zoom in/out or center figure (bottom right).

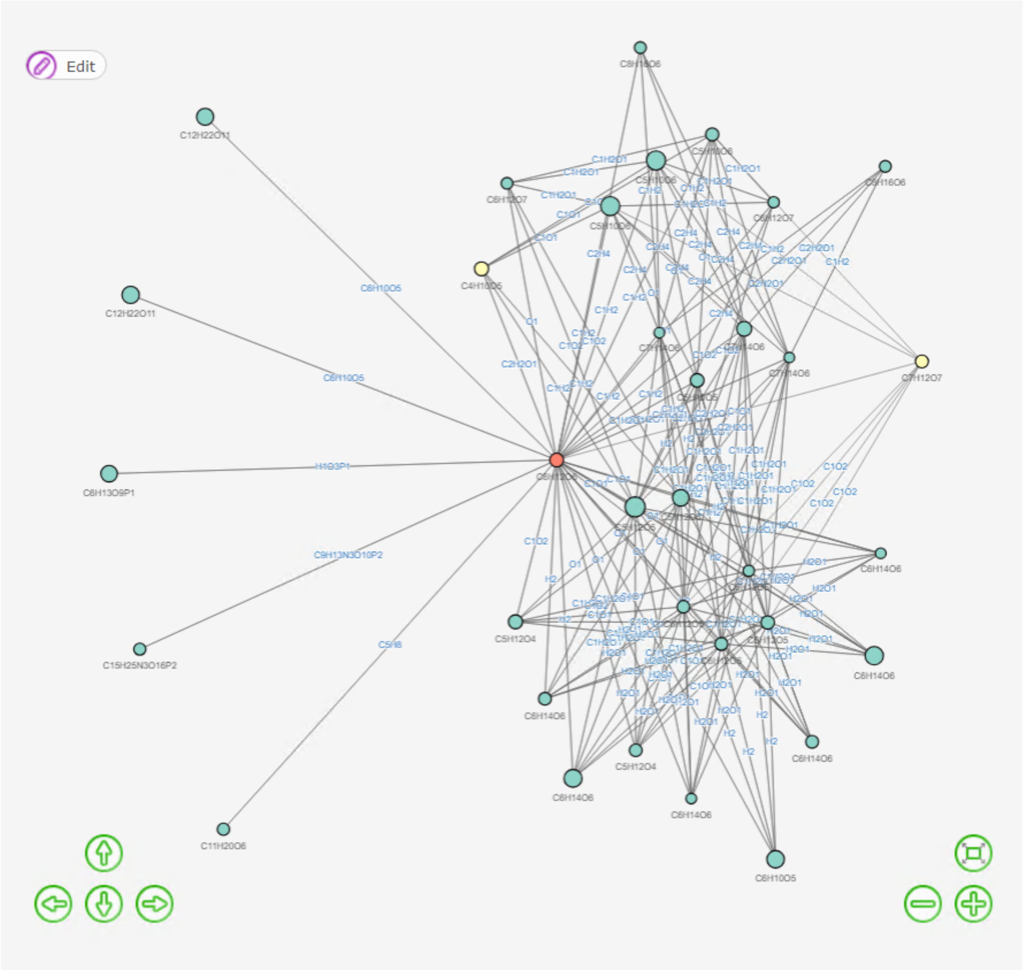


6. You can use the “Download plot” button to download a html webpage to visualize the network graph independent of the Shiny app, and the “Download csv” button to download the information of the nodes in the network. The download buttons will appear after hitting the plot button. Note: edits within the Shiny app will not go into the html file.

##### 3.2.4 Possible structures exploration

A figure + data table is provided to explore structures of the selected node in the network graph.

1. The figure shows the chemical structure of the annotated metabolites. If the node is annotated as a putative metabolite, only the known parts of the putative metabolite will be shown.
   Scroll left or right, or select the entry number, to visualize different annotations. Right click and select to save image.

2. In the data table, class has 3 possible entries: Metabolite if it is documented in database such as HMDB library; Putative metabolite if it is transformed from a metabolite through a biotransformation edge; and Artifact if it is transformed by an abiotic edge. Use the download button to download the data table

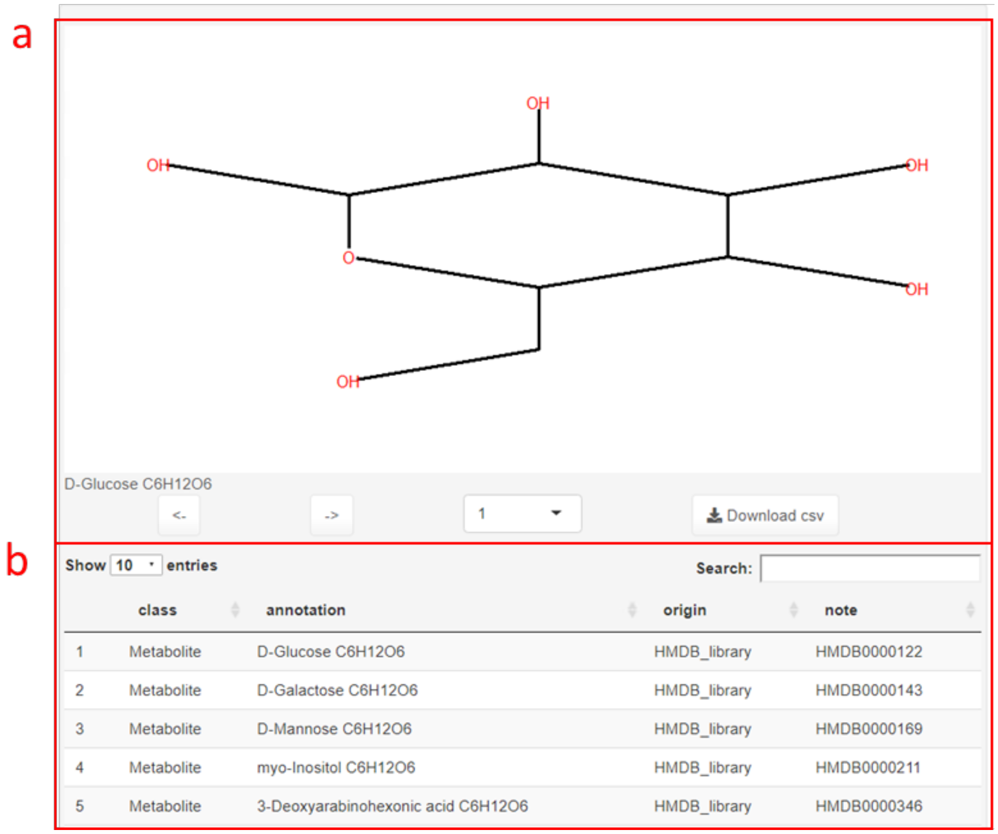


### 4 Troubleshooting

#### 4.1 Failing to install package `lc8`

Reinstall the packages `devtools` and `digest`.

#### 4.2 Cannot find `cplexAPI` even if the installation seems successful

Check **R** version used in **RStudio** to see if `cplexAPI` is installed under the same R version library. Which R library `cplexAPI` goes to depends on the R path specified in `Environment Variables`.

#### 4.3 Add **R** to `PATH`

1. Go to `Environment Variables`:
   search `PATH` in `windows` → open `edit Environment Variables` → `Environment Variables` or
   `control panel` → `system and security` → `System` → `Advanced system Settings` (on your left) → `Advanced` → `Environment Variables`

2. In the lower Panel select the `Path Variable` and select `Edit`, add the R path (`C:\Program Files\R\R-4.0.3\bin\x64`, if installed at default location) to the `Path Variable`.
3. You may need to restart computer for the R path to take effect.

#### 4.4 Add **Rtools40** to `PATH`

1. Add the path `C:\Rtools\bin` to the `Path Variable`in `Environment Variables`
2. Run the line in **R**:
   `writeLines('PATH="${RTOOLS40_HOME}\\usr\\bin;${PATH}"', con = "~/.Renviron")`
   Use the line below in R console to check for successfully adding Rtools40
   `Sys.which("make")`
   Expected output: `## "C:\\rtools40\\usr\\bin\\make.exe`

#### Supplementary Note 2 – NetID scoring

##### Scoring candidate node annotations

NetID scores every candidate node and edge annotation assigned in the candidate annotation step. The node scoring system aims to assign high scores to annotations that align observed ion peaks with known metabolites based on m/z, retention time, MS/MS, and/or isotope abundances.

Let the set of candidate annotations for node *u* be denoted as $\{a_1 \dots a_i \dots a_m\}$. For each node *u* and each of its candidate annotation $a_i$, let S(*u*, $a_i$) denotes the score of candidate annotation $a_i$ for node *u*. Different scoring components for candidate node annotations are defined as below:

(a) $\mathrm{S_{m/z}}(u, a_i)$ is negative when measured m/z differs from the calculated m/z of assigned molecular formula. A larger ppm difference between calculated formula m/z and measurement m/z results to lower scores. The default scale factor is -0.5. Let $a_{i,\mathrm{m/z}}$ be the calculated formula m/z of annotation $a_i$, and $u_{\mathrm{m/z}}$ be the measured m/z of node *u*, then

$$\mathrm{S_{m/z}}(u, a_i) = -0.5 \times \left| u_{\mathrm{m/z}} - a_{i,\mathrm{m/z}} \right| / \, u_{\mathrm{m/z}} \times 10^6 \quad (5)$$

(b) $\mathrm{S_{RT}}(u, a_i)$ is positive if the measured RT for the peak corresponding to node *u* matches to a known standard. A smaller difference between known and measured RT results in a higher score. Let $a_{i,\mathrm{RT}}$ is the known RT of annotation $a_i$, and $u_{\mathrm{RT}}$ be the measured RT of node *u*, then

$$\mathrm{S_{RT}}(u, a_i) = 1 - \left| u_{\mathrm{RT}} - a_{i,\mathrm{RT}} \right|, \text{ if } \left| u_{\mathrm{RT}} - a_{i,\mathrm{RT}} \right| < 0.5 \text{ min}$$

$$\text{Otherwise, } \mathrm{S_{RT}}(u, a_i) = 0 \quad (6)$$

(c) $\mathrm{S_{MS2}}(u, a_i)$ is positive if the measured MS2 spectrum of node *u* matches the database MS2 spectrum of annotation $a_i$. A dot product scoring function is used to score the MS2 spectra similarity[1,2]. The intensities of the fragment ions in the MS2 spectra are rescaled so that the highest fragment ion is set to 1. MS2 spectrum is represented as a data table containing m/z and corresponding relative intensity. Data tables for two spectra (one from experiment and one from database) are merged by m/z, which yields two equal-length vectors to represent relative intensity for experimental measured MS2 spectrum of $u$ ($W_u$) and database MS2 spectrum of $a_i$ ($W_{a_i}$). Dot product (DP) and score for MS2 match ($\mathrm{S_{MS2}}(u, a_i)$) are defined as below.

$$DP = \frac{\sum W_u W_{a_i}}{\sqrt{\sum W_u^2 \times \sum W_{a_i}^2}} \quad (7)$$

$$\mathrm{S_{MS2}}(u, a_i) = \mathrm{DP}, \text{ if } \mathrm{DP} > 0.5$$

$$\text{Otherwise } \mathrm{S_{MS2}}(u, a_i) = 0 \quad (8)$$

(d) $\mathrm{S_{database}}(u, a_i)$ is positive if the annotated formula $a_i$ exists in HMDB. We give a positive score to a primary seed node annotation if that annotated formula exists in HMDB.

$$\mathrm{S_{database}}(u, a_i) = 0.5, \text{ if } a_i \text{ in HMDB}$$

$$\text{Otherwise, } \mathrm{S_{database}}(u, a_i) = 0 \quad (9)$$

(e) $\mathrm{S_{missing\_isotope}}(u, a_i)$ is negative if an isotopic peak is missing. We penalize a formula annotation if it passes the intensity threshold (default at $5 \times 10^4$) but does not have isotopic peaks of specified elements.

The default isotope being evaluated is $^{37}Cl$. Any other elements, such as $^{13}C$ or $^{18}O$, can be included by users.

$$\mathrm{S}_{\mathrm{missing\_isotope}}(u, a_i) = -1, \text{ if isotopic peak is missing}$$
$$\text{Otherwise } \mathrm{S}_{\mathrm{missing\_isotope}}(u, a_i) = 0 \quad (10)$$

(f) $\mathrm{S}_{\mathrm{rule}}(u, a_i)$ is negative if annotation $a_i$ violates basic chemical rules. We strongly penalize formulae that violate basic chemical rules, including a negative RDBE (ring and double bond equivalents), and unlikely element ratios in metabolites (O/P < 3, O/Si < 2).

$$\mathrm{S}_{\mathrm{rule}}(u, a_i) = -10, \text{ if chemical rules are violated}$$
$$\text{Otherwise, } \mathrm{S}_{\mathrm{rule}}(u, a_i) = 0 \quad (11)$$

(g) $\mathrm{S}_{\mathrm{derivative}}(u, a_i)$ is a non-negative score that reflects annotation $a_i$ for node $u$ gains confidence that derived from its parent node *p* with candidate annotation *h*. This is particularly helpful in annotating abiotic peaks. For example, annotation of glutamate sodium adduct will be given a positive $\mathrm{S}_{\mathrm{derivative}}$ when its parent node is annotated as glutamate with high score $\mathrm{S}_{\mathrm{parent}}(p, h)$. $\mathrm{S}_{\mathrm{parent}}(p, h)$ is calculated by summing up scores in (a)-(f).

$$\mathrm{S}_{\mathrm{derivative}}(u, a_i) = \mathrm{S}_{\mathrm{parent}}(p, h) - 0.5, \text{ if } \mathrm{S}_{\mathrm{parent}}(p, h) > 0.5$$
$$\text{Otherwise, } \mathrm{S}_{\mathrm{derivative}}(u, a_i) = 0 \quad (12)$$
$$\mathrm{S}_{\mathrm{parent}}(p, h) = \mathrm{S}_{\mathrm{m/z}}(p, h) + \mathrm{S}_{\mathrm{RT}}(p, h) + \mathrm{S}_{\mathrm{MS2}}(p, h) + \mathrm{S}_{\mathrm{database}}(p, h) + \mathrm{S}_{\mathrm{missing\_isotope}}(p, h) + \mathrm{S}_{\mathrm{rule}}(p, h) \quad (13)$$

A final score S(*u*, $a_i$) for each candidate annotation $a_i$ of node *u* is calculated by summing scores in (a)-(g).

$$\mathrm{S}(u, a_i) = \mathrm{S}_{\mathrm{m/z}}(u, a_i) + \mathrm{S}_{\mathrm{RT}}(u, a_i) + \mathrm{S}_{\mathrm{MS2}}(u, a_i) + \mathrm{S}_{\mathrm{database}}(u, a_i) + \mathrm{S}_{\mathrm{missing\_isotope}}(u, a_i) + \mathrm{S}_{\mathrm{rule}}(u, a_i) + \mathrm{S}_{\mathrm{derivative}}(u, a_i) \quad (14)$$

Note that for each node *u*, no annotation is always a candidate option. The node score for this null annotation is 0 at default. It can also be set to a positive value to decrease false positive annotations or a negative value to promote increased annotation completeness (at the expense of more false positives).

**Scoring candidate edge annotations**

The edge scoring system aims to assign high scores to edge annotations that correctly capture biochemical connections between metabolites (based on MS2 spectra similarity) and abiotic connections between metabolites and their mass spectrometry phenomena derivatives, such as isotopes and adducts. Biochemical, isotope, and adduct edge annotations are the most common types. Other less common abiotic connection types are described in a subsequent section.

Suppose we consider two nodes *u* and *v* that are connected by an edge (*u*, *v*). For each pair of nodes *u* and *v* such that there is an edge (*u*, *v*), let the set of candidate formula for node *u* and *v* be denoted as $\{a_1 \dots a_i \dots a_m\}$ and $\{b_1 \dots b_j \dots b_n\}$, respectively, and let the set of candidate atom differences for edge (*u*, *v*) be $\{D_1 \dots D_k \dots D_l\}$. Let $\mathrm{S}(u, v, a_i, b_j, D_k)$ be the score of choosing candidate formula $a_i$ for node *u*, candidate formula $b_j$ for node *v* and candidate atom difference $D_k$ for edge (*u*, *v*). Note that $\mathrm{S}(u, v, a_i, b_j, D_k)$ is set to be 0 if atom difference $D_k$ does not represent the formula difference of $a_i$ and $b_j$.

$$S(u, v, a_i, b_j, D_k) = 0, \text{ if } |a_i - b_j| \neq D_k$$

Different scoring components for candidate edge annotations are defined as below:

(a) When node *u* and *v* have experimental measured MS2 spectra, $S_{\text{MS2\_similarity}}(u, v, a_i, b_j, D_k)$ is defined for a biochemical edge, and is a positive score if two connected nodes *u* and *v* have MS2 similarity. $S_{\text{MS2\_similarity}}$ is determined using the dot product (DP), as described in previous section, and reverse dot product (DP_R), which evaluates the neutral ion loss similarity in the MS2 spectra[1]. In calculating DP_R, data tables for two spectra (one from node u and one from node v) are merged by [precursor m/z – fragment m/z]. Two equal-length vectors representing the relative intensity of measured MS2 spectrum of $u$ and $v$ are denoted as $R_u$ and $R_v$ respectively. $S_{\text{MS2\_similarity}}$ is set at 0 for abiotic edges.

$$\text{DP} = \frac{\sum W_u W_v}{\sqrt{\sum W_u^2 \times \sum W_v^2}} \quad (15)$$

$$\text{DP\_R} = \frac{\sum R_u R_v}{\sqrt{\sum R_u^2 \times \sum R_v^2}} \quad (16)$$

$$S_{\text{MS2\_similarity}}(u, v, a_i, b_j, D_k) = \max(\text{DP}, \text{DP\_R}), \text{ if } \max(\text{DP}, \text{DP\_R}) > 0.3$$

$$\text{Otherwise, } S_{\text{MS2\_similarity}}(u, v, a_i, b_j, D_k) = 0 \quad (17)$$

(a) $S_{\text{co\_elution}}(u, v, a_i, b_j, D_k)$ is defined for an abiotic edge, and is a negative score if the RT of two connected nodes differ more than a threshold (0.05 min), given the formula difference of $a_i$ and $b_j$ matches the atom difference defined by $D_k$. $S_{\text{co\_elution}}$ is set at 0 for biochemical edges.

$$S_{\text{co\_elution}}(u, v, a_i, b_j, D_k) = -5 \times |u_{\text{RT}} - v_{\text{RT}}|, \text{ if } |u_{\text{RT}} - v_{\text{RT}}| \geq 0.05 \text{ min}$$

$$\text{Otherwise, } S_{\text{co\_elution}}(u, v, a_i, b_j, D_k) = 0 \quad (18)$$

(b) $S_{\text{type}}(u, v, a_i, b_j, D_k)$ is defined for all edges, given the formula difference of $a_i$ and $b_j$ matches the atom difference defined by $D_k$, and is a non-negative score depending on the connection type of edge, which is defined by $D_k$, including biotransformation, adduct, isotope and fragment (Supplementary Table 1, 2). The magnitude of scores reflects the empirical confidence in the annotation type when certain atom differences occur, and can be adjusted based on user preferences.

$$S_{\text{type}}(u, v, a_i, b_j, D_k) = 0, \text{ if } D_k \in \text{biotransformation}$$

$$S_{\text{type}}(u, v, a_i, b_j, D_k) = 0.5, \text{ if } D_k \in \text{adduct}$$

$$S_{\text{type}}(u, v, a_i, b_j, D_k) = 2, \text{ if } D_k \in \text{isotope}$$

$$S_{\text{type}}(u, v, a_i, b_j, D_k) = 0.3, \text{ if } D_k \in \text{common neutral loss} \quad (19)$$

(c) For each $D_k \in$ isotope, $S_{isotope\_intensity}(u, v, a_i, b_j, D_k)$ is defined for isotope edge (*u, v*) where $b_j$ is the isotopic derivative of $a_i$ with atom difference of $D_k$, and is a negative score if the measured isotope peaks deviate from expected natural abundance. The score for an isotope edge depends on how likely the ratio of measured and expected isotopic intensity ($\text{Ratio}_{\text{isotope}}$) is observed in an empirical normal distribution $\text{N}(1, \sigma_{\text{isotope}}^2)$. Isotopes of all elements included in the atom difference table are evaluated.

$$\text{Ratio}_{\text{isotope}} = \frac{v_{\text{intensity}} \,/\, u_{\text{intensity}}}{\text{Expected isotopic intensity ratio } (a_i, b_j, D_k)} \quad (20)$$

$$S_{isotope\_intensity}(u, v, a_i, b_j, D_k) = log_{10}\left[\frac{P\left(\mu = \mathrm{Ratio}_{\mathrm{isotope}} \middle| \mathrm{N}\left(1, \sigma_{\mathrm{isotope}}^2\right)\right)}{P\left(\mu = 1 \middle| \mathrm{N}\left(1, \sigma_{\mathrm{isotope}}^2\right)\right)}\right] \quad (21)$$

$\sigma_{\mathrm{isotope}}$ is empirically defined as below, so that when measured isotope intensity is close to detection limit, a larger $\sigma_{\mathrm{isotope}}$ (a widened distribution, which is more tolerant to discrepancy) will be used.

$$\sigma_{\mathrm{isotope}} = 0.2 + 10^{3 - log_{10}(v_{\mathrm{intensity}})} \quad (22)$$

**Less common edge annotations**

LC-MS metabolomics may include additional abiotic relationships. In orbitrap data, these include oligomers, multi-charge species, heterodimers, in-source fragments of known or unknown metabolites[3], and ringing artifact peaks surrounding high intensity ions[4,5]. These relationships were included in NetID as additional edge types, which are evaluated for all m/z pairs within a predefined RT range (0.2 min). Associated scores are provided at the end of the section.

(d) Oligomer and multi-charge species. An oligomer/multi-charge edge is assigned between two nodes *u* and *v*, if their m/z satisfy

$$|v_{\mathrm{m/z}} - \mathrm{n} \times u_{\mathrm{m/z}}| < u_{\mathrm{m/z}} \times 10\ \mathrm{ppm},\ \mathrm{n} \in \{\text{positive integers}\} \quad (23)$$

(e) Heterodimer. Heterodimer peak (node *v*) may be observed when one abundant metabolite (node *u*) forms ion cluster with other ion species (node *t*). We examine nodes that have intensity above $10^5$, and assign a heterodimer edge between two nodes *u* and *v* if their m/z difference satisfy

$$|(v_{\mathrm{m/z}} - u_{\mathrm{m/z}}) - t_{\mathrm{m/z}}| < u_{\mathrm{m/z}} \times 10\ \mathrm{ppm} \quad (24)$$

(f) In-source fragments. Such peaks may be observed when one abundant metabolite breaks up into fragments during the ionization process.

Database MS2 of known metabolites can be used to identify known ion fragment peaks[3]. If candidate annotation $b_j$ of node *v* is annotated with a HMDB ID associated with database MS2 spectrum, and m/z of node *u* matches to a fragment m/z in $b_j$'s MS2 spectrum, then a database fragment edge will connect such two nodes. That is,

$$u_{\mathrm{m/z}} \in \text{Database MS2 spectrum of candidate annotation } b_j \text{ of node } v \quad (25)$$

Measured MS2 spectra can also be used to identify fragment peaks (including covering unknowns not present in MS2 database). If node *v* is associated with a measured MS2 spectrum, and m/z of another node *u* matches to a fragment m/z in the MS2 spectra, then an experiment fragment edge will connect such two nodes. That is,

$$u_{\mathrm{m/z}} \in \text{Measured MS2 spectrum of node } v \quad (26)$$

(g) Ringing artifacts. Ringing peaks are artifact peaks (node *v*) often observed on both sides of the m/z of an intense ion peak (node *u*) in Fourier-transformed MS instrument including orbitrap. We examine nodes that have intensity above $10^6$, and assign a ringing artifact edge between two nodes if two nodes satisfy

$$50\ \mathrm{ppm} < |\ v_{\mathrm{m/z}} - u_{\mathrm{m/z}}\ |\ /\ u_{\mathrm{m/z}} < 1000\ \mathrm{ppm}$$

$$u_{\mathrm{intensity}}\ /\ v_{\mathrm{intensity}} > 50 \quad (27)$$

Scoring of these additional abiotic edges follow the same rules described in the "Scoring edge annotations" section with additional $S_{type}$ defined as below.

$$S_{type}(u, v, a_i, b_j, D_k) = 0.5, \text{ if } D_k \in \text{oligomer or multi-charge}$$
$$S_{type}(u, v, a_i, b_j, D_k) = 0, \text{ if } D_k \in \text{heterodimer}$$
$$S_{type}(u, v, a_i, b_j, D_k) = 0.3, \text{ if } D_k \in \text{database MS2 fragment}$$
$$S_{type}(u, v, a_i, b_j, D_k) = 1, \text{ if } D_k \in \text{measured MS2 fragment}$$
$$S_{type}(u, v, a_i, b_j, D_k) = 2, \text{ if } D_k \in \text{ringing artifacts} \quad (28)$$

A final edge annotation score $S(u, v, a_i, b_j, D_k)$ for choosing candidate formula $a_i$ for node *u*, candidate formula $b_j$ for node *v* and candidate atom difference $D_k$ for edge (*u*, *v*) is calculated by summing scores in (h)-(o).

$$S(u, v, a_i, b_j, D_k) = S_{MS2\_similarity}(u, v, a_i, b_j, D_k) + S_{co\_elution}(u, v, a_i, b_j, D_k) + S_{type}(u, v, a_i, b_j, D_k) + S_{isotope\_intensity}(u, v, a_i, b_j, D_k) \quad (29)$$

#### Supplementary Note 3. Pseudocode for NetID algorithm

The whole algorithm runs in the following workflow:

1. Input data and data cleaning
2. Initializing and defining NodeSet and EdgeSet
3. Expanding candidate annotation through edge propagation
4. Defining CplexSet
5. Scoring candidate node and edge annotations
6. Global optimization
7. Network annotation
8. Output

### Note: description following the “#” sign are comments, and will not be run by the code.

**1. Input data and data cleaning**

    # Input
    **Define** *Mset* as a list, read in
        Experimental MS1 data (containing mz, RT and intensity from LC-MS)
        Experimental MS2 data (associated with MS1 data)
        ionization mode
        HMDB library file
        HMDB library MS2 files (pos or neg)
        Known library file (with curated RT)
        Atom difference table (rule table for biotransformations, adducts, isotopes, etc.)

    # Remove background peaks and duplicated entries
    **For each** peak
        **IF** its intensity in procedure blank > 0.5-fold of that in biological samples
            **Remove** the peak
    **For** any two (or more) peaks,
        **IF** their mz difference is within *mz_tol* **AND** RT difference within *rt_tol*
            **Create** a new entry by merging multiple entries
                Take the median *mz* and *RT* of entries as new *mz* and *RT*
                Take the largest intensity value of all entries in each sample as sample intensity
            **Remove** old duplicated entry peaks

**2. Initializing and defining NodeSet and EdgeSet**

    # NodeSet: Each peak is a node, and becomes an entry in nodeset.
        **Define** *NodeSet* as a list,
        **For each** *node* in *NodeSet*,
            store one peak’s *mz*, *RT*, *intensity*, *MS2*

        # Set up seed nodes
        **For each** *node* in *NodeSet*,
            **IF** *mz_difference* < 10 ppm by comparing measured mz to all formulae in HMDB library

44 **Add** HMDB ID, formula and class information to corresponding node
45
46 # Adjust systematic measurement errors
47 **For all** nodes in *NodeSet* that has at least one HMDB entry,
48 **Linear regression** using measured mz values of selected nodes and their HMDB formula mz
49 an absolute mz adjustment factor $\varepsilon_{\text{absolute}}$ (independent of measured mz)
50 a relative mz adjustment factor $\varepsilon_{\text{relative}}$ (linearly dependent on measured mz)
51 **For each** node *u* in *NodeSet*
52 **Recalculate** measured mz by applying
53 $u_{\text{mz,adjusted}} = u_{\text{mz,measured}} \times (1 + \varepsilon_{\text{relative}}) + \varepsilon_{\text{absolute}}$
54
55 # EdgeSet: Each edge connects two nodes by a mass difference defined in atom difference table
56 **Define** *EdgeSet* as a list,
57 **For each** pair of node *u* and *v* (assuming $v_{\text{mz}} > u_{\text{mz}}$), and for each difference $D_i$ in atom difference table,
58 **IF** $|\,(v_{\text{m/z}} - u_{\text{m/z}}) - D_i\,| < v_{\text{m/z}} \times 10\ \text{ppm}$,
59 **IF** $D_i$ is a Biotransformation connection (defined in atom difference table)
60 **Add** an edge with *node1* = *u*, *node2* = *v*, and related info for $D_i$ to *EdgeSet*
61 **IF** $D_i$ is an Abiotic connection (defined in atom difference table) **AND**
62 **IF** $|\,v_{\text{RT}} - u_{\text{RT}}\,| < 0.2\ \text{min}$
63 **Add** an edge with *node1* = *u*, *node2* = *v*, and related info for $D_i$ to *EdgeSet*.
64
65 # EdgeSet expansion with additional abiotic connections (see manuscript methods)
66 # including oligomers, multi-charge species, heterodimers, in-source fragments, etc.
67 **For each** pair of node *u* and *v*
68 **IF** $|\,v_{\text{RT}} - u_{\text{RT}}\,| < 0.2\ \text{min}$ **AND**
69 **IF** properties of node u and v satisfy the criteria for additional abiotic connections
70 **Add** an edge with *node1* = *u*, *node2* = *v*, and related info for $D_i$ to *EdgeSet*.
71
72 **3. Expanding candidate node annotation through edge propagation**
73 # By applying the atom difference of edge (u, v) on the formula assigned to seed node u,
74 # we can derive a new candidate formula for the connected node v.
75 # Iterating the process to all candidate formulae of node u through edge (u, v) will
76 # further expand candidate formulae for node v.
77 # Seed nodes formulae from HMDB belong to Metabolite class.
78 **For each** *edge (u, v)* connecting node *u*, *v* in *EdgeSet* **AND** $D_i$ is a Biotransformation connection
79 **For each** candidate formula of node *u*, $u_{\text{formula}}$, that belongs to Metabolite class
80 **IF** calculated mz of $u_{\text{formula}} + D_{\text{i,formula}}$ is within 5 ppm of measured mz of node *v*
81 **Add** combined formula ($u_{\text{formula}} + D_{\text{i,formula}}$) with Metabolite class to node *v*
82 **For each** *edge (u, v)* connecting node *u*, *v* in *EdgeSet* **AND** $D_i$ is an Abiotic connection
83 **For each** candidate formula of node u, $u_{\text{formula}}$
84 **IF** calculated mz of $u_{\text{formula}} + D_{\text{i,formula}}$ is within 5 ppm of measured mz of node *v*
85 **Add** combined formula ($u_{\text{formula}} + D_{\text{i,formula}}$) with Artifact class to node *v*
86 **REPEAT** LINE 81-84 (above 4 lines) three times (three rounds of expansion via abiotic connections)
87 **REPEAT** LINE 77-85 (above 9 lines) two times (two rounds of expansion via biotransformation connections)

**4. Defining CplexSet**

### CplexSet defines the network structure for global network optimization
### Each node may contain zero, one or more than one candidate node annotations
### Each candidate node annotation defines an ilp_node in CplexSet (ilp means integer linear programing)
### We use *ilp_nodes* to score and record each candidate node annotation
### Similarly, we use *ilp_edges* to score and record each candidate edge annotation
**For each** node *u* in *NodeSet*,
  **For each** candidate node annotation $a_i$ in *u,*
    **IF** the combination of *node_id*, *formula* and *class* of $a_i$ is not in *ilp_nodes*
      **Add** the candidate annotation $(u, a_i)$ to *ilp_nodes*
**For each** edge *(u,v)* in *EdgeSet*,
  **For each** atom difference $D_k$
    **For each** candidate node annotation $a_i$ in *u*, and $b_j$ in *v*
      **IF** combined formula $(a_i + D_{\mathrm{k,formula}})$ == $b_j$
        **Add** the candidate edge annotation $(u, v, a_i, b_j, D_k)$ to *ilp_edges*

**5. Scoring candidate node and edge annotations**

### The scoring system is to assign high scores to annotations that effectively align the experimentally observed
### ion peaks with prior metabolomics knowledge.
### See manuscript method section for more details on score terms.
**For each** candidate node annotation $(u, a_i)$, its score $S(u, a_i)$ is the sum of
  $\mathrm{S}_{\mathrm{m/z}}(u, a_i)$ # based on m/z accuracy,
  $\mathrm{S}_{\mathrm{RT}}(u, a_i)$ # based on RT of measured peaks and known standards
  $\mathrm{S}_{\mathrm{MS2}}(u, a_i)$ # based on MS2 of measured peaks and database MS2
  $\mathrm{S}_{\mathrm{database}}(u, a_i)$ # based on if the annotation $a_i$ exists in HMDB
  $\mathrm{S}_{\mathrm{missing_{isotope}}}(u, a_i)$ # based on if the expected isotopic peak for $a_i$ is missing
  $\mathrm{S}_{\mathrm{rule}}(u, a_i)$ # based on if $a_i$ violates basic chemical rules
  $\mathrm{S}_{\mathrm{derivative}}(u, a_i)$ # based on if $a_i$ is derived from a parent peak with a high annotation score
**For each** candidate node annotation $(u, v, a_i, b_j, D_k)$, its score $S(u, v, a_i, b_j, D_k)$ is the sum of
  $\mathrm{S}_{\mathrm{MS2\_similarity}}(u, v, a_i, b_j, D_k)$ # based on similarity of measured MS2 spectra of node *u* and *v*
  $\mathrm{S}_{\mathrm{co\_elution}}(u, v, a_i, b_j, D_k)$ # based on RT difference of node *u* and *v*
  $\mathrm{S}_{\mathrm{type}}(u, v, a_i, b_j, D_k)$ # based on the connection type, defined by $D_k$ in the atom difference table
  $\mathrm{S}_{\mathrm{isotope\_intensity}}(u, v, a_i, b_j, D_k)$ # based on the intensity ratio and expected natural abundance

**6. Global optimization**

### The goal is to find annotations for each node so as to maximize the sum of the scores across the network
### under the constraints that each node is assigned a single annotation,
### and that the network annotation is consistent.
### An example optimization problem using CPLEX in R can be found at
### https://cran.r-project.org/web/packages/cplexAPI/vignettes/cplexAPI.pdf
**Define** *x* as a vector of binary number
  # if $x_i$ = 1, the candidate node or edge annotation is selected in the global optimal network
  # if $x_i$ = 0, then the annotation is not selected.

Length(*x*) = (number of candidate node annotation) + (number of candidate edge annotation)
**Define** *Obj* as a vector with the same length of *x*
### *Obj* records score for each candidate node or edge annotation
### *Obj* · *x* is the total scores of the network. Global optimization maximizes the total scores.
*Obj* = c(scores for ilp_nodes, scores for ilp_edges)

### Constraints are defined as below.
### For a sample constraint $a_1x_1 + a_2x_2 \le b$,
### $a_1x_1 + a_2x_2$ is the left-hand side, b is the right-hand side and "≤" is the sense of the constraint
### $[a_1, a_2]$ is the constraint matrix, $[a_1, a_2] \cdot x$ is the left-hand side of a constraint
**Define** *mat* as a matrix
### *mat* · *x* is the left-hand side of the constraint.
### *mat* is a sparse matrix as most number in *mat* are zero.
Column number of *mat* = Length(*x*)
Row number of *mat* = Number of constraints
**Define** *triplet_mat* as a matrix
### we use triplet (*i,j,v*), i.e. the value (*v)* in the *i*th row, and *j*th column, to describe *mat*.
Column number of *triplet_mat* = 3
Row number of *triplet_mat* = number of non-zero entry in *mat*
**Define** *rhs* as a numeric vector
### *rhs* is the right-hand side of a constraint
Length(*rhs*) = Number of constraints
**Define** *sense* as a character vector
### *rhs* describes the signs between left- and right-hand sides
### Signs includes less or equal (L), equal (E), greater or equal (G)
Length(*sense*) = Number of constraints

### How constraint matrix is filled up is described below.
### (I) Constrain each peak has single annotation.
### Total number for this constraint = number of peaks
### for all annotation $a_i$ of peak *u*, sum ($x_{ai}$) = 1
**For each** candidate node annotation in *ilp_nodes*,
**Add** *i* = *peak_id*, *j* = *ilp_node_id*, *v* = 1 to *triplet_mat*
**Add** rep(1, number of peaks) to *rhs*
**Add** rep('E', number of peaks) to *sense*

### (II) Constrain each edge annotation exists only if related candidate node annotations exist.
### Total number for this constraint = number of candidate edge annotations * 2
### In candidate edge annotation $e(u,v,a_i,b_j,D_k)$, $x_e - x_{ai} \le 0$ and $x_e - x_{bj} \le 0$
$i_{current}$ = total number of constraints from (I)
**For each** candidate edge annotation in *ilp_edges*,
$i_{current} = i_{current} + 1$
**Add** $i = i_{current}$, *j* = *ilp_edge_id,* and *v* = 1;
$i = i_{current}$, *j* = *ilp_node_id* for $a_i$, and *v* = -1 to *triplet_mat*

$i_{current} = i_{current} + 1$
**Add** $i = i_{current}$, $j$ = *ilp_edge_id,* and $v = 1$;
$i = i_{current}$, $j$ = *ilp_node_id* for $b_j$, and $v = -1$ to *triplet_mat*
**Add** rep(0, number of candidate edge annotations * 2) to *rhs*
**Add** rep('L', number of candidate edge annotations * 2) to *sense*

### (III) Constrain an isotope annotation exists only if the isotope connection exists
### Total number for this constraint = number of candidate edge annotation that is an isotope connection
### In candidate edge annotation $e(u,v,a_i,b_j,D_k)$, assuming $b_j$ is an isotope annotation, $x_e - x_{bj} = 0$
$i_{current}$ = total number of constraints from (I-II)
**For each** candidate edge annotation in *ilp_edges* that is an isotope connection
$i_{current} = i_{current} + 1$
**Add** $i = i_{current}$, $j$ = *ilp_edge_id,* and $v = 1$;
$i = i_{current}$, $j$ = *ilp_node_id* for $b_j$, and $v = -1$ to *triplet_mat*
**Add** rep(0, number of candidate edge annotation that is an isotope connection) to *rhs*
**Add** rep('E', number of candidate edge annotation that is an isotope connection) to *sense*

### (IV) Constrain only one edge can exist between two nodes
### Total number for this constraint = number of multiple-edge events * 2
### When multiple edges exist between two nodes, we call it multiple-edge event
### Assuming candidate edge annotation $e(u,v,a_i,b_j,D_k)$, $e'(u,v,a_i,b_j,D_k)$ and multiple edges exist
### At most one edge exist: $x_e + x_{e'} + \ldots - x_{ai} \le 0$, $x_e + x_{e'} + \ldots - x_{bj} \le 0$
$i_{current}$ = total number of constraints from (I-III)
**For each** multiple edge event
$i_{current} = i_{current} + 1$
**Add** $i = i_{current}$, $j$ = *ilp_node_id* for $a_i$, and $v = -1$ to *triplet_mat*
**For each** candidate annotation $e$ that exist between node u and node v with $a_i$ and $b_j$ annotation
**Add** $i = i_{current}$, $j$ = *ilp_edge_id* for $e$, and $v = 1$ to *triplet_mat*
$i_{current} = i_{current} + 1$
**Add** $i = i_{current}$, $j$ = *ilp_node_id* for $b_j$, and $v = -1$ to *triplet_mat*
**For each** candidate annotation $e$ that exist between node u and node v with $a_i$ and $b_j$ annotation
**Add** $i = i_{current}$, $j$ = *ilp_edge_id* for $e$, and $v = 1$ to *triplet_mat*
**Add** rep(0, number of multiple edge event * 2) to *rhs*
**Add** rep('L', number of multiple edge event * 2) to *sense*

### Pass parameters to CPLEX optimization
**Add** *CPLEX_para* = list(*nc* = Length(*x*), *nr* = number of constraint # number of columns and rows
*CPX_MAX*, # indicating maximization will be performed
*obj*, *rhs*, *sense*, # described above
*cnt*, *ind*, *val*, # describing mat in compressed sparse column (CSC) format
*lb* = 0, *ub* = 1, *ctype* = "B" # x's lower and upper bound, and its type is binary
) to *CplexSet*

### CPLEX optimization

```
        # ilp_solution contains a vector of binary number that
        # denotes if a candidate node or edge annotation is selected for the global optimal network.
        ilp_solution = Run_cplex(CplexSet)
        optimized_nodes = Filter ilp_nodes that selected in ilp_solution
        optimized_edges = Filter ilp_edges that selected in ilp_solution
```

**7. Network annotation**

```
    # Seeds are node annotations that have direct annotations from HMDB,
        Define optimized_seed_nodes = Filter optimized_nodes that have HMDB annotations
        Define optimized_nodes_M = Filter optimized_nodes that are Metabolite class annotation
        Define optimized_nodes_A = Filter optimized_nodes that are Artifact class annotation
        Define optimized_edges_M = Filter optimized_edges that are Biotransformation connections
        Define optimized_edges_A = Filter optimized_edges that are Abiotic connections

    # The output network is an overlay of a biotransformation network and an abiotic network
        Define g_bio, g_abiotic, g_all as graphs,
            g_bio = graph (edges = optimized_edges_M,
                           nodes = optimized_nodes_M)
            g_abiotic = graph (edges = optimized_edges_A,
                               nodes = optimized_nodes that exist in optimized_edges_A)
            g_all = g_bio + g_abiotic

    # Every node annotation in the network can trace back to seed annotation
        Define bio_dist as a distance matrix,
            # distance in row i and column j records
            # the shortest distance from node i in optimized_seed_nodes to node j in optimized_nodes_M
            bio_dist = shortest.paths(graph = g_bio,
                                      from = optimized_seed_nodes,
                                        to = optimized_nodes_M)
        Define abiotic_dist as a distance matrix,
            # distance in row i and column j records
            # the shortest distance from node i in optimized_nodes_M to node j in optimized_nodes_A
            abiotic_dist = shortest.paths(graph = g_abiotic,
                                          from = optimized_nodes_M,
                                            to = optimized_nodes_A)

    # Path annotations to nodes
        For each node M in optimized_nodes_M
            Find seed node H that has shortest distances to M among all optimized_seed_nodes in bio_dist
            Define path as the intermediate edges and nodes connecting from H to M
            Add path annotation = c(HMDB name of H,
                                    HMDB formula of H,
                                    # Atom differences are specified by edge annotations in path
                                    1st step atom difference, "->", intermediate node formula,
```

…
last step atom difference, “->”, Formula of *M*) to node *M*
### for acetyl-thiamine peak: “thiamine C12H16N4O1S1 + C2H2O1 -> C14H18N4O2S1”
**For each** node *A* in optimized_nodes_A
**Find** Metabolite class node *M* that has shortest distances to *A* among all in *abiotic_dist*
**Define** *path* as the intermediate edges and nodes connecting from *M* to *A*
**Add** *path annotation* = c(Formula of *M*,
### Atom differences are specified by edge annotations in path
1st step atom difference, “->”, intermediate node formula,
…
last step atom difference, “->”, Formula of *A*) to node *A*
### for glutamate sodium acetate adduct peak:
# “C5H9N1O4 + Na1H-1 -> C5H8Na1N1O4 + C2H4O2 -> C7H12N1Na1O6”

**8. Output**
### csv format
**For all** peaks,
compiles peak_id, medMz, medRt, log10_inten, class, formula, ppm_error, path annotation
**Exports** as NetID_output.csv
### Shiny R visualization
**Save** all information as NetID_output.RData for Shiny R visualization
